# Identification of RBPMS as a smooth muscle master splicing regulator via proximity of its gene with super-enhancers

**DOI:** 10.1101/563304

**Authors:** Erick E. Nakagaki-Silva, Clare Gooding, Miriam Llorian, Aishwarya Griselda Jacob, Frederick Richards, Adrian Buckroyd, Sanjay Sinha, Christopher W.J. Smith

## Abstract

Alternative splicing (AS) programs are primarily controlled by regulatory RNA binding proteins (RBPs). It has been proposed that a small number of master splicing regulators might control cell-specific splicing networks and that these RBPs could be identified by proximity of their genes to transcriptional super-enhancers. Using this approach we identified RBPMS as a critical splicing regulator in differentiated vascular smooth muscle cells (SMCs). RBPMS is highly down-regulated during phenotypic switching of SMCs from a contractile to a motile and proliferative phenotype and is responsible for 20% of the AS changes during this transition. RBPMS directly regulates AS of numerous components of the actin cytoskeleton and focal adhesion machineries whose activity is critical for SMC function in both phenotypes. RBPMS also regulates splicing of other splicing, post-transcriptional and transcription regulators including the key SMC transcription factor Myocardin, thereby matching many of the criteria of a master regulator of AS in SMCs.

## Introduction

Alternative splicing (AS) is an important component of regulated gene expression programmes during cell development and differentiation, usually focusing on different sets of genes than transcriptional control (Blencowe, 2006). AS programs re-wire protein-protein interaction networks (Buljan et al., 2012; J. D. Ellis et al., 2012; Yang et al., 2016), as well as allowing quantitative regulation by generating mRNA isoforms that are differentially regulated by translation or mRNA decay (McGlincy & Smith, 2008; Mockenhaupt & Makeyev, 2015). Coordinated cell-specific splicing programs are determined by a combination of *cis*-acting transcript features and *trans*-acting factors that compose “splicing codes” (Barash et al., 2010; Chen & Manley, 2009; X. D. Fu & Ares, 2014). The interactions between the *trans* component RNA binding proteins (RBPs) and the *cis* component regulatory elements in target RNAs coordinate the activation and repression of specific splicing events. Many regulatory proteins, including members of the SR and hnRNP protein families, are quite widely expressed, while others are expressed in a narrower range of cell types (David & Manley, 2008; X. D. Fu & Ares, 2014). A further conceptual development of combinatorial models for splicing regulation has been the suggestion that a subset of RBPs act as master regulators of cell-type specific AS networks (Jangi & Sharp, 2014). The criteria expected of such master regulators include that: i) they are essential for cell-type specification or maintenance, ii) their direct and indirect targets are important for cell-type function, iii) they are likely to regulate the activity of other splicing regulators, iv) they exhibit a wide dynamic range of activity, which is not limited by autoregulation, and v) they are regulated externally from the splicing network, for example by transcriptional control or post-translational modifications. It was further suggested that expression of such splicing master regulators would be driven by transcriptional super-enhancers, providing a possible route to their identification (Jangi & Sharp, 2014). Super-enhancers are extended clusters of enhancers that are more cell-type specific than classical enhancers and that drive expression of genes that are essential for cell-type identity, including key transcription factors (Hnisz et al., 2013). By extension, RBPs whose expression is driven by super-enhancers are expected to be critical for cell-type identity and might include master regulators of tissue-specific AS networks (Jangi & Sharp, 2014).

Vascular smooth muscle cells (SMCs) are important in cardiovascular physiology and pathology (Bennett, Sinha, & Owens, 2016; Fisher, 2010; Owens, Kumar, & Wamhoff, 2004). Unlike skeletal and cardiac muscle SMCs exhibit phenotypic plasticity and are not terminally differentiated (Owens et al., 2004) (Fig. 1A). In healthy arteries, vascular SMCs exist in a differentiated contractile state. In response to injury or disease, the SMC phenotype switches towards a more synthetically active, motile and proliferative state (Fisher, 2010; Owens et al., 2004). The transcriptional control of SMC phenotypic switching has been intensely studied, but the role of post-transcriptional regulation has been relatively neglected (Fisher, 2010). For example, some markers of the contractile state, such as h-Caldesmon and meta-Vinculin, arise via AS (Owens et al., 2004), but nothing is known about the regulation of these events. A number of known splicing regulators, including PTBP1, CELF, MBNL, QKI, TRA2B, and SRSF1, have been implicated in the regulation of individual SMC-specific ASEs, but these proteins are not restricted to differentiated SMCs and most act primarily in the de-differentiated state (Gooding et al., 2013; Gooding, Roberts, & Smith, 1998; Gromak, Matlin, Cooper, & Smith, 2003; Shukla & Fisher, 2008; van der Veer et al., 2013; Xie et al., 2017). Indeed, global profiling confirmed a widespread role of PTBP1 in repressing exons that are used in differentiated mouse aorta SMCs (Llorian et al., 2016), but did not identify RBPs that act as direct regulators of the differentiated state. Biochemical identification of such RBPs is hampered by the fact that SMCs rapidly dedifferentiate in cell culture conditions.

**Figure 1.**
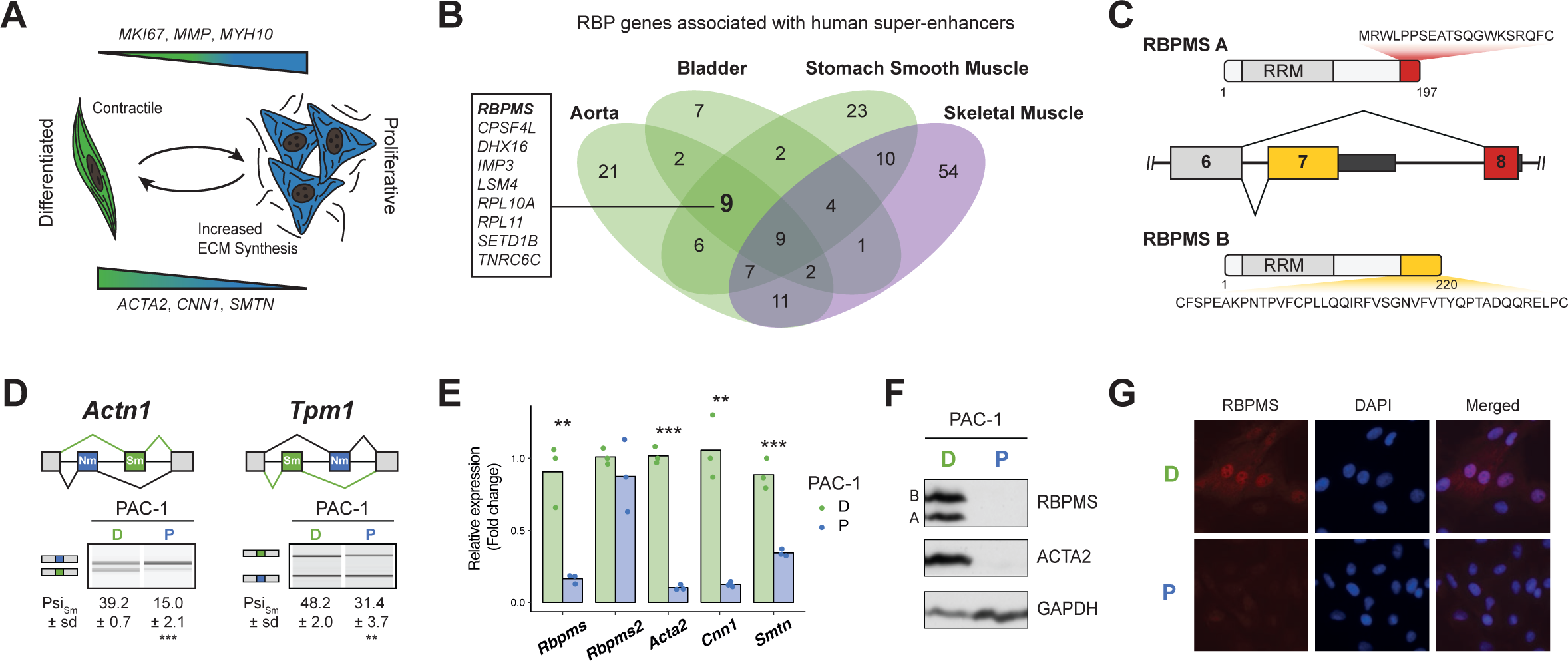
RBPMS is associated with SMC super-enhancers and is highly expressed in the differentiated PAC1 cells. (**A**) Diagram of the SMC dedifferentiation. SMC markers of differentiation and dedifferentiation are respectively shown at the bottom and top of the diagram. (**B**) Venn diagram of RBP genes associated with super-enhancers across different human smooth muscle tissues. Skeletal muscle was used as an outlier. RBPs common to all smooth muscle tissues but not skeletal muscle are shown on the left. (**C**) Schematic of the AS event determining the two major RBPMS isoforms, RBPMS A (red) and RBPMS B (yellow). (**D**) RT-PCR analysis of SMC splicing markers, *Actn1* and *Tpm1*, in differentiated (D) and proliferative (P) PAC1 cells. Schematic of the regulated mutually exclusive splicing events on top and respective isoforms products on the left. Values shown are the quantified Psi (percent spliced in) of the smooth muscle isoforms (Sm). (**E**) qRT-PCR analysis of *Rbpms* (all isoforms), *Rbpms2* and SMC differentiation markers *Acta2, Cnn1* and *Smtn*, in PAC1 cells D (green) and P (blue). Expression was normalized to the average of two housekeepers (*Gapdh* and *Rpl32*) and the mean of the relative expression is shown (*n*=3). Statistical significance was performed using Student’s t-test (* P < 0.05, ** P < 0.01,*** P < 0.001). (**F**) Western blots for RBPMS in D and P PAC1 cells. ACTA2 is a SMC differentiation marker and GAPDH a loading control. A and B indicates the two RBPMS isoforms. (**G**) Immunofluorescence in D and P PAC1 cells for RBPMS. DAPI staining for nuclei.

Here, we used the approach suggested by Jangi and Sharp, to identify candidate AS master regulators as RBP-encoding genes whose cell-specific expression is driven by super-enhancers (Jangi & Sharp, 2014). We identified <u>R</u>NA <u>B</u>inding <u>P</u>rotein with <u>M</u>ultiple <u>S</u>plicing (RBPMS), a protein not previously known to regulate splicing, as a critical regulator of numerous AS events in SMCs. RBPMS is highly expressed in differentiated SMCs, where it promotes AS of genes that are important for SMC function. These include many components of the actin cytoskeleton and focal adhesion machineries, modulation of whose function is key to the transition from contractile to motile phenotypes. RBPMS also targets other splicing regulators, post-transcriptional regulators and the key SMC transcription factor Myocardin where RBPMS promotes inclusion of an exon that is essential for maximal SMC-specific activity. RBPMS therefore meets many of the criteria expected of an AS master regulator in SMCs, and its identification validates the approach of identifying key cell-specific regulators via the super-enhancer-proximity of their genes.

## Results

### RBPMS is a SMC splicing regulator

To identify potential SMC master splicing factors we used a catalog of 1542 human RBPs (Gerstberger, Hafner, & Tuschl, 2014) and data-sets of super-enhancers from three human SMC-rich tissues (aorta, bladder, stomach smooth muscle) and skeletal muscle (Hnisz et al., 2013)(Supplementary File 1). Nine RBP genes were associated with super-enhancers in all SMC tissues but not skeletal muscle (Fig 1B). Using a set of super-enhancer-associated genes in the dbSUPER database (Khan & Zhang, 2016), we identified two candidates, of which only RBPMS was shared with our original 9 candidates. Examination of *RBPMS* expression in human tissues from the Genotype-Tissue Expression (GTEX) project (Consortium, 2013) showed the top 8 expressing tissues to be SMC rich, including three arteries (Fig. S1A). RNA-Seq data from rat aorta SMCs showed that *Rbpms* levels decreased 3.8 fold during de-differentiation from tissue to passage 9 cell culture (Fig. S1B-D), in parallel with known SMC transcriptional markers (*Acta2, Cnn1, Smtn*, (Fig. S1C,D) and AS events (*Tpm1* and *Actn1*, Fig. S1E,F). Moreover, of the starting candidate RBPs (Fig 1B) *Rbpms* was the most highly expressed of those that were down-regulated from tissue to culture (Fig. S1C green labels). Expression of the *Rbpms2* paralog also decreased upon de-differentiation, but its absolute level of expression was >10-fold lower than *Rbpms* (Fig. S1C,D). Other RBPs implicated in AS regulation in SMCs (PTBP1, MBNL1, QKI) either showed more modest changes or higher expression in de-differentiated cells (Fig. S1C,D). In the PAC1 rat SMC line *Rbpms* mRNA levels decreased by ∼10-fold between differentiated and proliferative states in parallel with SMC marker AS events (Fig. 1D) and genes (Fig. 1E). *Rbpms2* was expressed at much lower levels than *Rbpms*, and did not alter expression between PAC1 cell states (Fig 1E). RBPMS protein decreased to undetectable levels in proliferative PAC1 cells in parallel with smooth muscle actin (ACTA2) (Fig 1F). Immunofluorescence microscopy also showed higher levels of RBPMS in differentiated PAC1 cells where it was predominantly nuclear (Fig 1G), consistent with the hypothesis that it regulates splicing.

To further investigate RBPMS expression we cloned cDNAs from PAC1 cells, representing seven distinct mRNA isoforms. These encoded two major protein isoforms (RBPMSA and RBPMSB) sharing a common N-terminus and RNA Recognition Motif (RRM) domain. RBPMSA and B differed by short C-terminal tails encoded by alternative 3’ end exon 7 and exon 8 (Fig. 1C), and corresponded in size to the two protein bands seen in western blots (Fig. 1F). Other mRNA isoforms differed by inclusion or skipping of exon 6 and by alternative 3’ UTR exon inclusion. RBPMS and RBPMS2, which are 70% identical, have a single RRM domain that is responsible for both RNA binding and dimerization (Sagnol et al., 2014; Teplova, Farazi, Tuschl, & Patel, 2016). Optimal RBPMS binding sites consist of tandem CACs separated by a spacer of ∼1-12 nt (Farazi et al., 2014; Soufari & Mackereth, 2017). We found significant enrichment of CACN_1-12_CAC motifs within and upstream of exons that are less included in differentiated compared to cultured rat aorta SMCs (Fig. S1H). These are locations at which many splicing regulators mediate exon skipping (X. D. Fu & Ares, 2014; Witten & Ule, 2011). Consistent with this, the SMC-specific mutually exclusive exon pairs in *Actn1* and *Tpm1* (Gooding & Smith, 2008; Southby, Gooding, & Smith, 1999) both have conserved clusters of CAC motifs upstream of the exon that is skipped in differentiated SMCs (see below).

In summary, the presence of super-enhancers at the *RBPMS* gene in SMC-rich tissues, its wide dynamic range of expression between differentiated SMCs and other tissues and proliferative SMCs (Fig. 1, S1), the nuclear localization of RBPMS, and the presence of potential RBPMS binding sites adjacent to known SMC-regulated exons are all consistent with the hypothesis that RBPMS might act as a master regulator of AS in differentiated SMCs.

### RBPMS promotes a differentiated SMC splicing program

To investigate the roles of RBPMS in shaping SMC transcriptomes we manipulated levels of RBPMS expression in differentiated and proliferative PAC1 cells (Fig. 2A,B). We used siRNAs to knockdown all *Rbpms* isoforms in differentiated PAC1 cells, achieving ∼75% depletion with no effect on the SMC marker ACTA2 (Fig. 2B). In parallel, proliferative PAC1 cells were transduced with pINDUCER lentiviral vectors (Meerbrey et al., 2011) to allow Doxycycline inducible over-expression of RBPMSA. No basal RBPMS expression was observed in proliferative cells, but upon induction substantial expression was observed from the FLAG-RBPMSA vector but not from the empty lentiviral vector (LV) (Fig. 2B). The effects of manipulating RBP levels can sometimes be compensated by related family members (Mockenhaupt & Makeyev, 2015). However, *Rbpms2* levels were not affected by any of the treatments and *Rbpms2* knockdown, either alone or in combination with *Rbpms*, had no effects upon tested AS events (data not shown).

**Figure 2.**
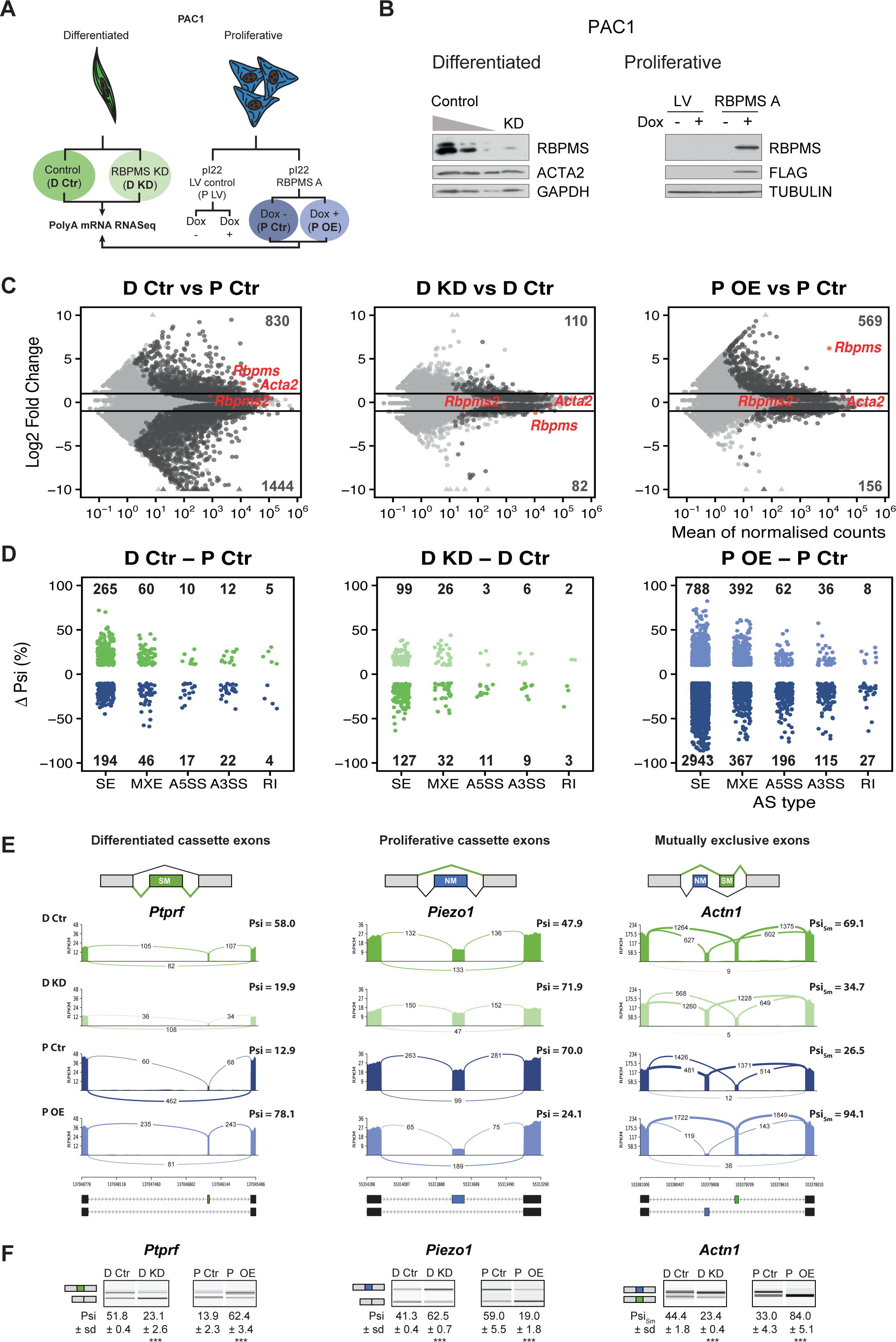
RBPMS regulates AS in PAC1 cells. (**A**) Schematic of experimental design of RBPMS knockdown and overexpression in PAC1 cells. (**B**) Western blots for RBPMS in PAC1 knockdown, left, and inducible lentiviral overexpression, right. FLAG antibodies were also used for the overexpression of 3xFLAG tagged RBPMS. Antibody targeting ACTA2 as a SMC differentiation marker and GAPDH and TUBULIN as loading controls. (**C**) MA plots of alterations in mRNA abundance in PAC1 dedifferentiation, left, RBPMS knockdown, middle, and RBPMS A overexpression, right. Dark gray: genes with significant changes (p-adj < 0.05). Light gray: genes with p-adj ≥ 0.05. Red: *Rbpms, Rbpms2* and the SMC marker, *Acta2*. Numbers of up and down-regulated are shown at top and bottom. Horizontal lines; log2 fold change = 1 and −1. (**D**) AS changes (FDR < 0.05 and ΔPsi greater than 10%) in PAC1 cell dedifferentiation, left, RBPMS knockdown, middle, and RBPMS A overexpression, right. ASE were classified into skipped exon (SE), mutually exclusive exon (MXE), alternative 5′ and 3′ splice site (A5SS and A3SS) and retained intron (RI) by rMATS. Numbers indicate the number of significant ASE of each event type between the conditions compared. (**E**) Sashimi plots of selected ASEs. *Ptprf* is shown as a differentiated cassette exon (green), *Piezo1* as a proliferative cassette exons (blue) and *Actn1* as a MXE. The numbers on the arches indicate the number of reads mapping to the exon-exon junctions. Psi values for the ASE are indicated for each condition. Values correspond to the mean Psi calculated by rMATS. In the case of the *Actn1* MXE, the Sm exon percent of inclusion is shown (Psi_Sm_). Schematic of the mRNA isoforms generated by the alternative splicing are found at the bottom as well as the chromosome coordinates. (**F**) RT-PCR validation of ASEs from panel E. Values shown are the mean of the Psi ± standard deviation (*n* = 3). Statistical significance was calculated using Student’s t-test (* P < 0.05, ** P < 0.01,*** P < 0.001). See Fig S2 for more ASEs validated in the RBPMS knockdown and RBPMS A overexpression by RT-PCR.

RNA samples from *Rbpms* knockdown and overexpression experiments were prepared for Illumina poly(A) RNAseq. Data was analysed for changes in mRNA abundance using DESEq2 (Love, Huber, & Anders, 2014) (Fig. 2C, Supplementary File 2), and for changes in AS using rMATS (Shen et al., 2014) (Fig. 2D, Supplementary File 3). In addition to analysis of effects of RBPMS depletion and overexpression, comparison of the differentiated and proliferative PAC1 control samples revealed changes associated with differentiation state. Principal Component Analysis based on mRNA abundance showed clear separation of differentiated and proliferative samples (PC1, 82% variance, Fig. S2A). This suggests that at the level of mRNA abundance the differences between differentiated and proliferative PAC1 cells far outweigh any effects of manipulating RBPMS levels. Consistent with this, RBPMS knockdown was associated with ∼10-fold fewer changes at the transcript abundance level (110 increased, 82 decreased, padj < 0.05, fold change > 2-fold) compared to the differentiated vs proliferative control comparison (830 increased, 1444 decreased, Fig. 2C). RBPMS overexpression led to an intermediate number of changes in mRNA abundance levels, but only 29 genes were affected by both overexpression and knockdown, and of these only 4 genes other than *Rbpms* were regulated reciprocally.

RBPMS knockdown and overexpression had substantial effects at the level of AS affecting all types of AS event (Figs. 2D, S2B). RBPMS knockdown led to changes in 318 AS events (FDR < 0.05, |ΔPSI| > 0.1, where PSI is <u>P</u>ercent <u>S</u>pliced In), which was only 2-fold less than the number of events regulated between control differentiated and proliferative cells. Cassette exons were the largest group of events, with roughly equal numbers of up-and down-regulated exons (Fig. 2D). RBPMS overexpression led to a larger number of AS changes (4934 regulated events), probably resulting from the combination of RBPMS expression in excess of levels usually present in differentiated PAC1 cells (Fig. 2C) and also because we expressed the more active RBPMSA isoform (see below). Cassette exons affected by RBPMS overexpression were strongly skewed (80%) towards greater exon skipping. A subset of ASEs observed in the RNA-Seq experiments, encompassing RBPMS activated and repressed cassette exons and mutually exclusive exons were validated by RT-PCR (Fig. 2E,F, Fig. S2E,F). ΔPSI values determined by RT-PCR and RNA-Seq were in good agreement (Fig. S2G). As an additional negative control, doxycycline induction of empty lentiviral transduced cells had no effect on RBPMS regulated AS events (Fig. S2D).

In each of the three comparisons, there was little overlap between genes regulated at the levels of splicing and mRNA abundance (Fig. S2C). However, there were substantial overlaps of ASEs regulated by RBPMS knockdown, overexpression and PAC1 phenotype (Fig 3A). Twenty percent of ASEs regulated in PAC1 cell differentiation were congruently regulated by RBPMS knockdown, representing 40% of ASEs affected by RBPMS knockdown (Fig 3A, 3B left panel). The high correlation (R^2^= 0.95) suggests that for these 127 events changes in RBPMS expression are sufficient to explain their differentiation-specific splicing changes. Similarly RBPMS overexpression shared 180 events in common with PAC1 differentiation status (28% of differentiation specific events). The correlation between ΔPSI for AS events co-regulated in PAC1 differentiation and RBPMS overexpression (R^2^= 0.86) was lower than for RBPMS knockdown and represented only 3.7% of events regulated by RBPMS overexpression (Fig 3A, 3B right panel). Hierarchical clustering of cassette exons regulated between PAC1 phenotypes across all 12 samples also revealed two clusters of RBPMS-responsive events where knockdown and overexpression were sufficient to reciprocally convert the splicing pattern to that of the other cellular phenotype (Fig. 3E, clusters 1 and 4, containing RBPMS activated and repressed exons respectively).

**Figure 3.**
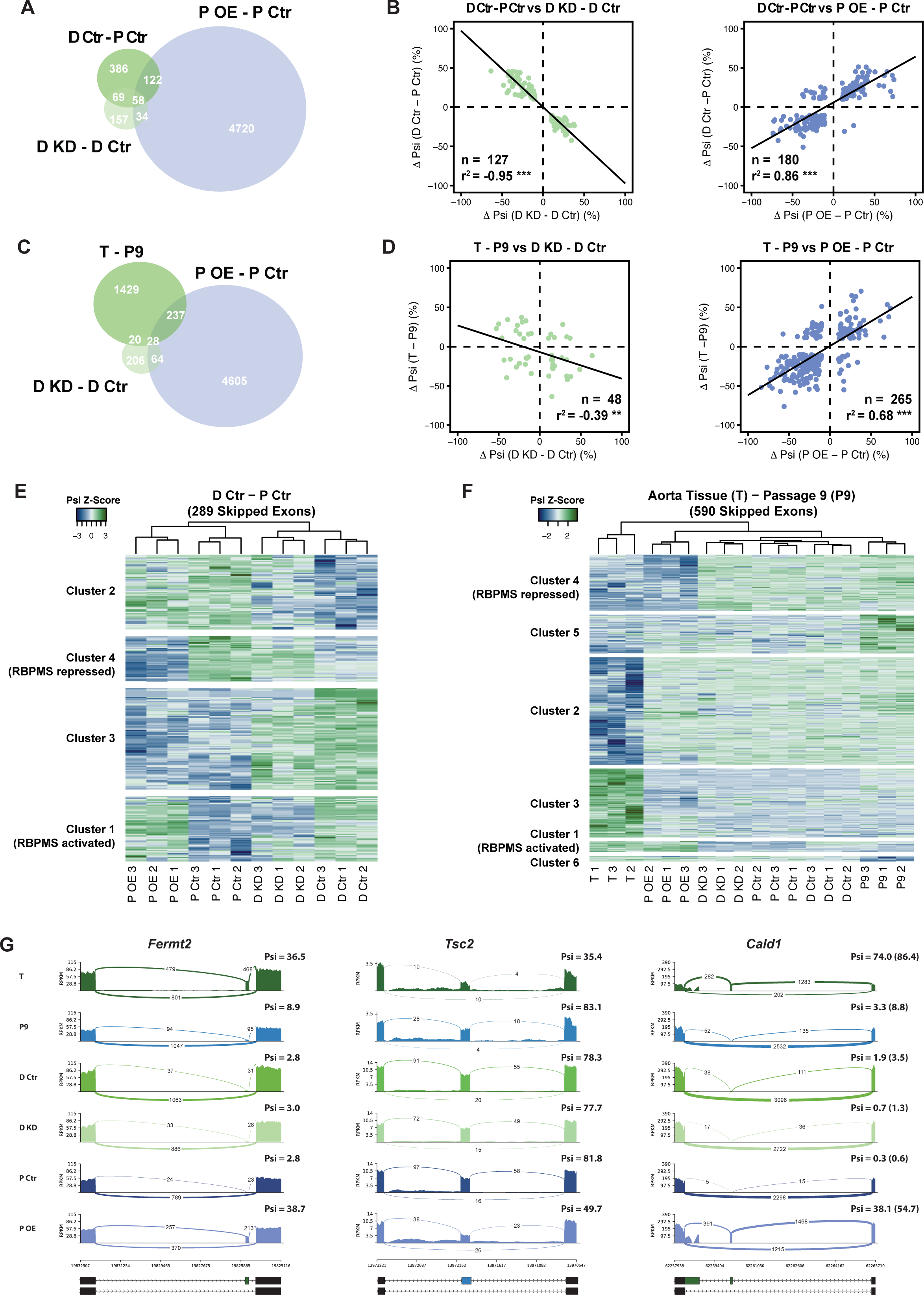
RBPMS recapitulates the AS program of differentiated PAC1 and aorta tissue. (**A**) Venn diagram of significant ASEs (FDR < 0.05 and ΔPsi cutoff of 10%) identified in the PAC1 dedifferentiation, RBPMS knockdown and RBPMS A overexpression comparisons. (**B**) ΔPsi correlation of overlapping ASEs (FDR < 0.05 and ΔPsi cutoff of 10%) from the PAC1 cells dedifferentiation and RBPMS knockdown, left, or RBPMS A overexpression, right. The line indicates the linear regression model. Statistical significance was carried out by Pearson correlation test. n is the number of ASEs assessed, r^2^ is the correlation coefficient and the p-value indicates the statistical significance of the correlation. (**C**) Venn diagram of significant ASEs identified in RBPMS knockdown, overexpression and aorta tissue to passage 9 comparisons. (**D**) ΔPsi correlation of overlapping ASEs from the rat aorta tissue dedifferentiation and RBPMS knockdown, left, or RBPMS A overexpression, right. (**E**) Heatmap of 289 SEs regulated in the PAC1 dedifferentiation comparison (D Ctr - P Ctr). Each column represents a replicate sample (1-3) from RBPMS knockdown (D Ctr and D KD) or overexpression (P Ctr and P OE). Rows are significant SE events from PAC1 dedifferentiation. Rows and columns were grouped by hierarchical clustering. Blue and green colors in the Z-score scaled rows represent low and high Psi values. (**F**) Heatmap of 590 SEs regulated in the rat aorta tissue dedifferentiation comparison (T - P9). Each column represents one sample from either aorta tissue dedifferentiation (T and P9), RBPMS knockdown (D Ctr and D KD) or RBPMS overexpression (P Ctr and P OE). Rows are significant SE events from the T - P9 comparison. Rows and columns were grouped by hierarchical clustering. Blue and green colors in the Z-score scaled rows represent low and high Psi values. (**G**) Sashimi plots of selected ASEs highly regulated in the aorta tissue and RBPMS overexpression. *Fermt2* is an RBPMS activated exon from Cluster 1 in panel F. *Tsc2* is an RBPMS repressed exon from Cluster 4 of panel E. *Cald1* has an RBPMS activated exon 4 and downstream 5 splice site on exon 3b. For Cald1, a manually calculated Psi in parentheses takes into account the A5SS which was not included in the rMATS annotation.

Many AS events in differentiated PAC1 cells do not reach the fully differentiated splicing pattern characteristic of tissue SMCs. However, for some events overexpression of RBPMS led to splicing patterns similar to tissue SMCs. For example the *Actn1* SM exon is included to 69% in differentiated PAC1 cells, but to 93-94% in RBPMS overexpressing cells (Fig 2E) and aorta tissue (Fig. S1F). We therefore hypothesized that some ASEs regulated by RBPMS overexpression but not knock-down, might reflect tissue SMC AS patterns that are not usually observed in cultured SMCs. To address this possibility we used RNA-Seq data monitoring de-differentiation of rat aorta SMCs (Fig. S1B,C). Of 1714 ASES regulated between tissue and passage 9 SMCs, 265 (15%) were also regulated by RBPMS overexpression in PAC1 cells (Fig. 3C,D, R^2^= 0.68). Strikingly, hierarchical clustering of cassette exons regulated between aorta tissue and passage 9 cultured SMCs showed that the RBPMS overexpression sample clustered together with tissue, away from all other samples (Fig. 3F). Two clusters of AS events shared very similar splicing patterns in tissue and RBPMSA overexpression, and differed in all remaining samples (Fig. 3F). Cluster 1 comprised 28 RBPMS-activated exons, of which 20 were not regulated between PAC1 phenotypes. For example, inclusion of a cassette exon in *Fermt2* was observed only in tissue and RBPMS overexpression samples (Fig. 3G). Similarly, inclusion of *Cald1* exon 4 in conjunction with a downstream 5’ splice site on exon 3a, producing the hCald1 marker in tissue SMCs (Fig 3G, S3C), and inclusion of the meta-vinculin exon (Fig. S3A) were also only seen upon RBPMS overexpression. Cluster 4 contained exons that are skipped in tissue and upon RBPMS overexpression (e.g. *Tsc2*, Fig. 3G), the majority of which are not regulated in PAC1 differentiated cells. Likewise *Tpm1* mutually exclusive exon 3 is skipped nearly completely upon RBPMS overexpression in a similar pattern to tissue (Fig. S3B). Upregulation of RBPMS expression therefore appears to be sufficient to promote a subset of splicing patterns usually only observed in differentiated SMCs *in vivo*.

### RBPMS directly regulates exons with associated CAC motifs

To address whether RBPMS directly regulates target exons we looked for enrichment of its binding motif (Farazi et al., 2014) adjacent to cassette exons regulated by RBPMS knockdown or overexpression. CACN_1-12_CAC motifs were significantly enriched around exons that were activated or repressed by RBPMS, with a similar position-dependent activity as other splicing regulators (Fig 4A). Exons repressed by RBPMS showed strong enrichment of motifs within the exon and the immediate ∼80 nt upstream intron flank, while exons activated by RBPMS showed motif enrichment within the downstream intron flank. Consistent with the contribution of RBPMS to the AS changes between PAC1 differentiation states, CACN_1-12_CAC motifs were also enriched upstream of and within exons that are more skipped in differentiated cells, and downstream of exons that are more included in differentiated cells (Fig 4A). Moreover, in the set of exons activated by RBPMS overexpression, CACN_1-12_CAC motifs were not only enriched downstream, but also significantly depleted in the repressive locations within and upstream of the exon (Fig. 4A,B). This suggests that binding in repressive locations might be dominant over activation.

**Figure 4.**
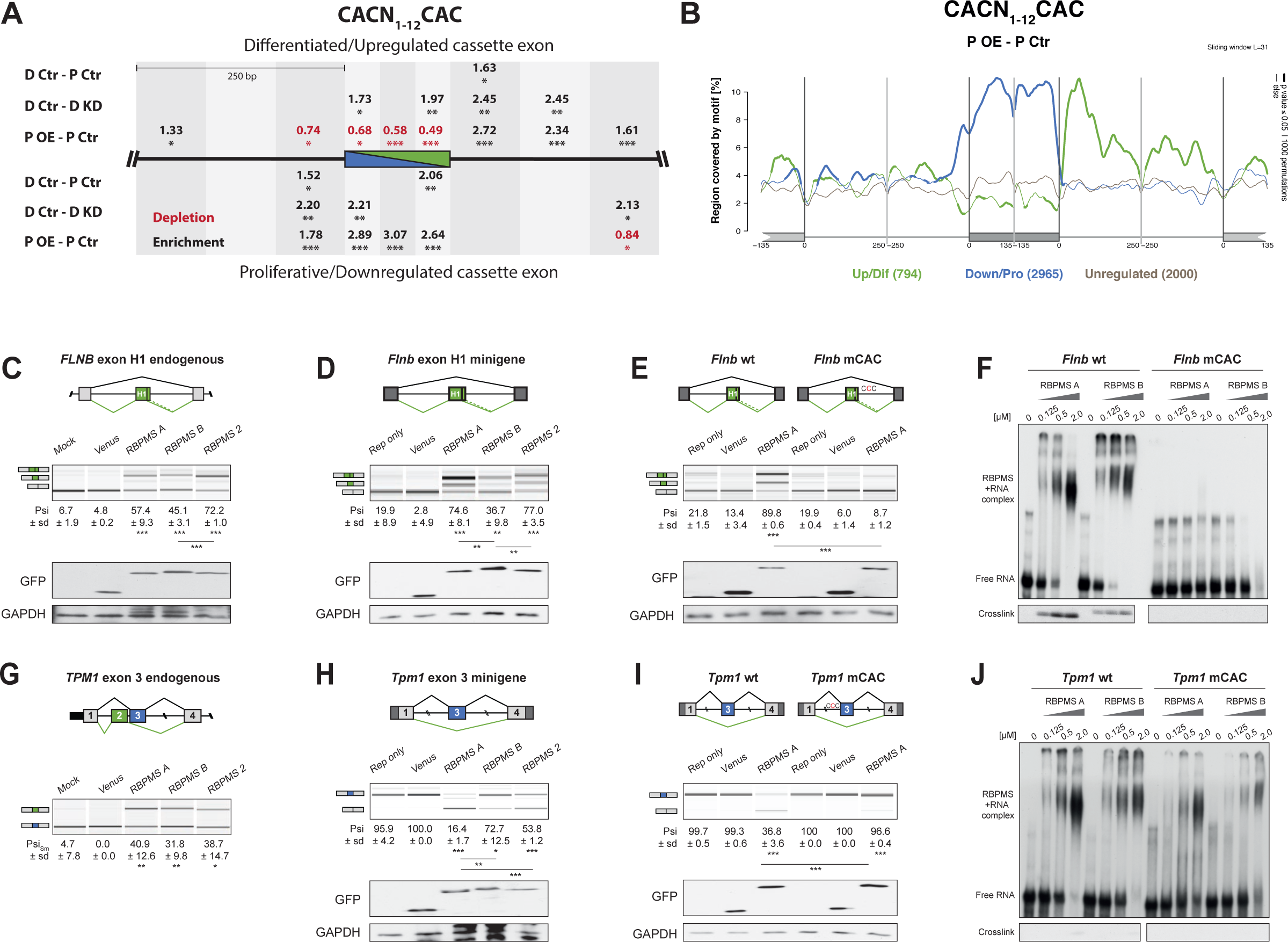
RBPMS directly regulates exons associated with CAC motifs. (**A**) RBPMS motif (CACN_1-12_CAC) enrichment in differentially alternatively spliced cassette exons in the PAC1 dedifferentiation (D Ctr - P Ctr), RBPMS knockdown (D Ctr - D KD) and RBPMS overexpression (P OE - P Ctr) comparisons. Values indicate degree of motif enrichment (black) or depletion (red). Statistical significance: * P < 0.05, ** P < 0.01,*** P < 0.001). (**B**) RBPMS motif (CACN_1-12_CAC) map from RBPMS overexpression (P OE - P Ctr). Upregulated (green), downregulated (blue) and unregulated (grey) cassette exons are shown. Statistical significance is indicated by the line width. (**C**) RBPMS overexpression in HEK293T cells and RT-PCR of endogenous *FLNB* exon H1 splicing. Schematic of *Flnb* exon H1 as a differentiated cassette exon activated by RBPMS. The splicing pattern of *FLNB* was tested upon overexpression of Venus tagged RBPMS A, B and 2. A mock and a Venus control were also tested in parallel. (**D**) Effect of RBPMS overexpression on rat *Flnb* splicing reporter in HEK293T cells. Schematic of the *Flnb* reporter and RT-PCR for the splicing patterns of *Flnb* reporter upon coexpression of RBPMS isoforms and RBPMS2. (**E**) *Flnb* reporter with point mutations disrupting RBPMS motifs (CAC to CCC) were tested for RBPMS A regulation. Left wild-type reporter and right mutant CAC reporter (mCAC) RT-PCRs. (**F**) Electric mobility shift assay (EMSA) for *in vitro* binding of recombinant RBPMS A and B to *in vitro* transcribed wild-type and mCAC RNAs of *Flnb* exon H1 downstream intron sequence (top left and right). Lower panel: UV-crosslinking of same samples. (**G**) RBPMS overexpression in HEK293T cells and RT-PCR of endogenous *TPM1* MXE exon 2 and 3 splicing. The splicing pattern of *TPM1* was tested upon overexpression of Venus tagged RBPMS A, B and 2. A mock and a Venus control were also tested in parallel. (**H**) Effect of RBPMS overexpression on rat *Tpm1* splicing reporter in HEK293T cells. Schematic of the *Tpm1* reporter and RT-PCR for the splicing patterns of *Tpm1* reporter upon RBPMS isoforms and paralog. (**I**) *Tpm1* reporter with point mutations disrupting RBPMS motifs (CAC to CCC) were tested for RBPMS A regulation. Left wild-type reporter and right mutant CAC reporter (mCAC) RT-PCRs. (**J**) *In vitro* binding of recombinant RBPMS A and B to *in vitro* transcribed wild-type and mCAC RNAs of *Tpm1* exon 3 upstream intron sequence. EMSA, top and UV crosslinking, bottom. In (**F**) and (**J**), *in vitro* binding assays were carried out using recombinant protein in a serial dilution (1:4) in a range of 0.125 to 2 μM. In panels **C**-**E** and **G**-**I**, values are the mean Psi ± SD (*n* = 3). For *Flnb* cassette exon, the Psi is the sum of both short and long isoforms generated by a A5SS event. For *Tpm1* MXE, the Sm exon Psi is shown (Psi_Sm_). Schematics of the splicing isoforms indicate the PCR products, differentiated (green) and proliferative (blue). Statistical significance was calculated using Student’s t-test (* P < 0.05, ** P < 0.01,*** P < 0.001). Western blot anti-GFP and GAPDH, loading control, were carried out to assess RBPMS isoforms overexpression in HEK293. The western blot in (**C**) is also a representative of (**G**), since both RT-PCRs (*FLNB* and *TPM1*) are from the same overexpression experiment.

To test whether RBPMS regulates AS events by directly binding to CAC motifs we co-transfected HEK293 cells with RBPMS expression vectors and minigenes of representative activated (*Flnb*) and repressed (*Tpm1, Actn1*) exons, with potential RBPMS binding sites in expected locations for activation or repression (Fig. S4C-E). We initially established that transient expression of RBPMSA in HEK293 cells was sufficient to switch AS of endogenous *FLNB, TPM1, MPRIP* and *ACTN1* towards the SMC splicing pattern (Fig 4C,G, Fig. S4A). For comparison, we also transfected expression constructs for the RBPMSB isoform and the paralog RBPMS2. We found that RBPMSB had lower activity for some events (*ACTN1*, Fig. S4A), but that transfected RBPMS2 had similar activity to RBPMSA in all cases. RBPMSA and RBPMS2 also strongly activated inclusion of the *Flnb* H1 exon in a minigene context while RBPMSB activated to a lower extent (Fig 4D). *Tpm1* exon 3 is the regulated member of a pair of mutually exclusive exons and is repressed in SMCs (P. D. Ellis, Smith, & Kemp, 2004; Gooding, Roberts, Moreau, Nadal-Ginard, & Smith, 1994). MBNL and PTBP proteins promote this repression but are not sufficient to switch splicing (Gooding et al., 2013; Gooding et al., 1998). In contrast, RBPMSA expression was sufficient to cause a near complete switch from exon inclusion to skipping (Fig 4H). RBPMS2 had lower activity, but RBPMSB was by far the least active protein. Likewise, RBPMSA and RBPMS2 completely switched splicing of *Actn1* constructs (Gromak et al., 2003; Southby et al., 1999) from the NM to the SM mutually exclusive exon, while RBPMSB was nearly inactive (Fig. S4B). A construct containing only the *Actn1* SM exon was unresponsive to cotransfection, while a construct containing only the NM exon, which has 3 upstream CAC clusters (Fig. S4C) showed a complete switch from inclusion to skipping upon cotransfection of RBPMSA or RBPMS2, but not RBPMSB. Thus, for two mutually exclusive events RBPMSA is able to switch splicing to the SMC pattern by repressing the exon that is usually used in non-SMCs, while RBPMSB is less active.

We mutated CAC motifs in suspected binding sites to CCC, which disrupts RBPMS binding (Farazi et al., 2014). Mutation of 12 CACs downstream of the *FlnB* H1 exon had no effect on basal splicing, but the H1 exon was completely resistant to activation by RBPMSA (Fig 4E). Likewise, mutation of 9 CAC motifs upstream of *Tpm1* exon 3 had no effect on exon inclusion in the absence of RBPMSA, but completely prevented exon skipping in response to RBPMSA (Fig 4I, Fig. S4E), while mutations of individual clusters had intermediate effects (Fig. S4E). The response of both *Flnb* H1 exon and *Tpm1* exon 3 therefore depends on nearby CAC motifs. To test whether these are binding sites for RBPMS, we used *in vitro* transcribed RNAs and recombinant RBPMSA and B (Fig. 4F, J, Fig. S4F,G). Using both electrophoretic mobility shift assay (EMSA) and UV crosslinking, both RBPMSA and B were found to bind to the *FlnB* wild type RNA, but not to the mutant RNA (Fig. 4F). With *Tpm1*, RBPMSA and B bound to the WT RNA as indicated by EMSA assays (Fig. 4J). Binding was reduced by the mutations that abrogated RBPMSA repression of Tpm1 exon 3. These data therefore show that RBPMS can inhibit splicing by binding to sites upstream of exons (*Tpm1*) and activate splicing by downstream binding (*Flnb*).

### RBPMS regulated splicing targets the cytoskeleton and cell adhesion

To investigate the functional importance of RBPMS regulated AS we carried out Gene Ontology (GO) analysis of the genes whose splicing or expression levels were regulated by RBPMS (Fig5A, Fig. S5A, Tables S4, S5), or between SMC differentiation states (Fig. S5B,C). Splicing events affected by RBPMS-knockdown affected genes involved in processes, components and functions important for SMC biology, such as cytoskeleton, cell projection, cell junction organization and GTPase regulation (Fig 5A). These categories were very similar to those affected by AS during PAC1 cell dedifferentiation (Fig. S5B). Events regulated by RBPMS knockdown were also enriched within genes that are associated with super-enhancers in SMC tissues (Fig. 5B) and therefore inferred to be important for SMC identity. In contrast, the relevance of GO terms associated with RBPMSA overexpression to SMC biology was less clear (Fig. S5A), and overexpression AS targets were not enriched for aorta super-enhancer associated genes (p = 0.37). Likewise, at the RNA abundance level enriched GO terms associated with RBPMS knockdown or overexpression did not align with those associated with differentiation, mainly being associated with stress responses (Supplementary File 5).

**Figure 5.**
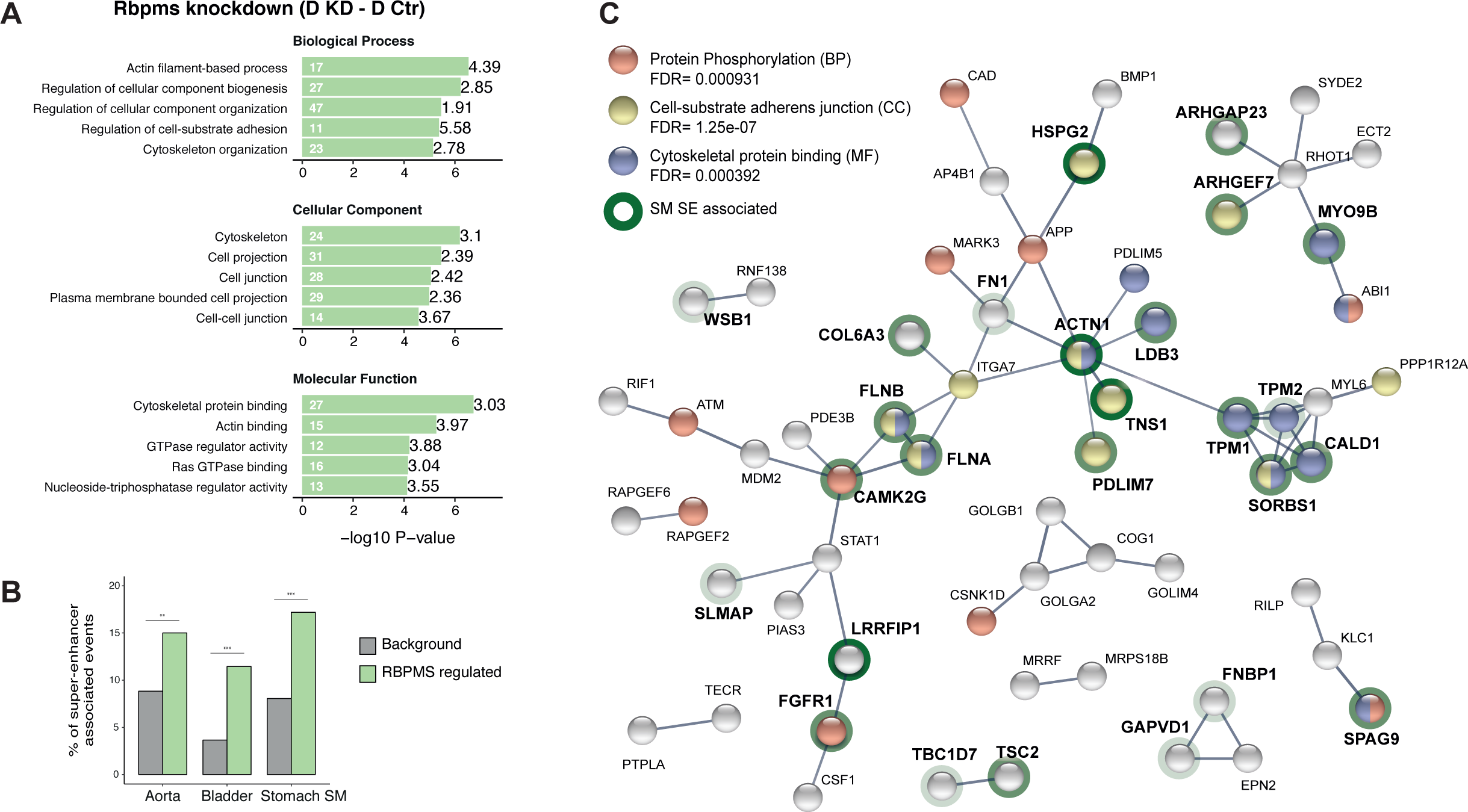
RBPMS regulates functionally important targets in SMCs. (**A**) GO analysis of genes with cassette exons regulated in RBPMS knockdown. The top five enriched GO terms in the three categories (biological process, molecular function and cellular component) are shown. Values within and in front of the bars indicate the number of genes in the enriched term and the enrichment relative to the background list. (**B**) Enrichment of exons regulated by RBPMS knockdown within genes associated with super-enhancers in smooth muscle tissues. Background set is all cassette exon events (regulated and unregulated) detected by rMATS in the same experiment. Significance determined by hypergeometic P-value. (**C**) PPI network of genes differentially spliced upon RBPMS knockdown and overexpression that overlap with the PAC1 and aorta tissue dedifferentiation datasets. PPI network was generated in STRING using experiments and database as the sources of interactions. Network edges represent the interaction confidence. Enriched GO terms (BP, biological process, MF, molecular function and CC, cellular component) were also included in the analysis and are indicated in red, blue and yellow. Super-enhancer associated gene names are in bold and are highlighted gray, light green or dark green shading according to whether they were super-enhancer associated in 1, 2 or 3 SMC tissues.

To further explore the consequences of RBPMS regulation we carried out network analysis using STRING (Szklarczyk et al., 2017) (Fig. 5C). We used only target AS events that are coregulated by RBPMS knockdown and PAC1 differentiation state (Fig. 3A,E) or by RBPMS overexpression and in aorta tissue (Fig. 3C,F), and restricted the output network to high confidence interactions. RBPMS targets comprised a network (Fig. 5C) focused on functions associated with cell-substrate adhesion (yellow nodes) and the actin cytoskeleton (blue). Six of the network proteins (SORBS1, CALD1, PDLIM5, PDLIM7, ACTN1 and ARHGEF7) are also components of the consensus integrin adhesome (Horton et al., 2015), which mechanically connects, and mediates signalling between, the actin cytoskeleton and the extracellular matrix. Underlining the importance of this network to SMC function, many of the genes are themselves super-enhancer-associated in one or more smooth muscle tissues (bold text, Fig. 5C). Actomyosin activity, and its mechanical connection to the extracellular matrix are important for both the contractile and motile states of SMCs (Min et al., 2012). It appears that RBPMS plays an important role in modulating the activity of this protein network to suit the needs of the two cell states. This control of a functionally coherent set of targets that are important for SMC function is consistent with the hypothesis that RBPMS is a master regulator of AS in SMCs.

### RBPMS controls post-transcriptional regulators

Among the direct targets of master splicing regulators are expected to be events controlling the activity of other AS regulators leading to further indirect AS changes. Consistent with this, we noted that RBPMS affected splicing of *Mbnl1* and *Mbnl2*. RBPMS promoted skipping of the 36 nt and 95 nt alternative exons of *Mbnl1* (here referred to as exons 7 and 8), as indicated by both knockdown and overexpression (Fig. 6A,B). In *Mbnl2* the 36 nt exon was fully skipped in all conditions but the 95 nt exon was repressed by RBPMS (Fig. 6C). Consistent with the RNA-Seq data, we obtained *Mbnl1* and *Mbnl2* cDNAs from PAC1 cells that varied by inclusion of the 36 and 95 nt exons. Corresponding shifts in MBNL1, but not MBNL2, protein isoforms could be observed by western blot (Fig 6D). These events affect the unstructured C-termini of MBNL proteins and have been shown to affect their splicing activity (Sznajder et al., 2016; Tabaglio et al., 2018; Tran et al., 2011), suggesting that RBPMS might indirectly affect some AS events by modulating MBNL activity. To test this, we made expression constructs of rat MBNL1 isoforms with and without exons 7 and 8 (FL, Δ7, Δ8, Δ7Δ8) and MBNL2 with and without exon 8 (FL and Δ8), obtained as cDNAs from PAC1 cells. When transfected into HEK293 cells, full length MBNL1 and 2 caused a shift from use of the downstream to an upstream 5’ splice site (5’SS) on *NCOR2* exon 47 (Fig 6E), an event that is differentially regulated by MBNL isoforms (Sznajder et al., 2016; Tran et al., 2011). The shorter MBNL isoforms showed lower activity in shifting towards the upstream 5’SS, although the difference was not statistically significant for the MBNL1 shorter isoforms (Fig. 6E). In proliferative PAC1 cells, *NCOR2* exon 47 mainly uses the upstream 5’SS, and knockdown of MBNL1 and 2 caused a significant shift to the downstream 5’SS (Fig. 6F). Overexpression of RBPMSA, also caused a small shift to the downstream 5’SS (Fig. 6F, also detected by rMATs, ΔPSI = 13%, FDR = 2 × 10^−6^), but had no effect when MBNL1 and MBNL2 were knocked down. These results are consistent with MBNL1 and 2 being direct regulators of Ncor2 splicing, with RBPMS acting indirectly by promoting production of less active MBNL isoforms.

**Figure 6.**
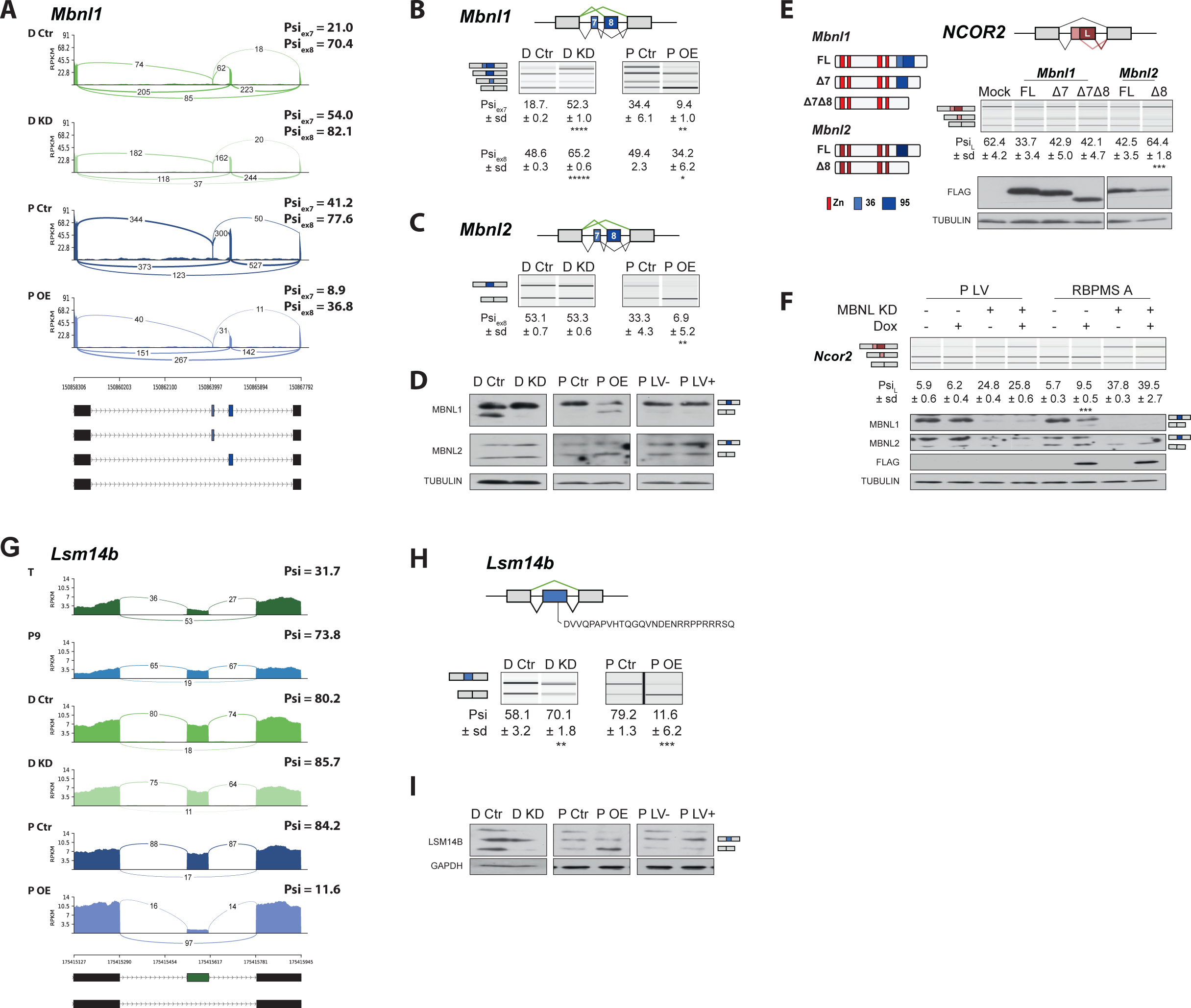
RBPMS regulates splicing of post-transcriptional regulators. (**A**) Sashimi plot of *Mbnl1* regulated exons. The numbers on top of the arches indicate the number of reads mapping to the exon-exon junctions. Mean of the Psi values calculated for each exon (exon 7 – 36 bp and exon 8 – 95 bp) are indicated for each condition (Psi_ex7_ and Psi_ex8_). Schematic of the different mRNA isoforms are found at the bottom. (**B**) RT-PCR validation of Mbnl1 exons 7 and 8 and (**C**) Mbnl2. Values shown are the mean of the Psi ± standard deviation (*n* = 3). The Psi values for each exon of Mbnl1 are shown. For *Mbnl2*, exon 7 isoforms were not detected in the RT-PCRs. Schematics of the splicing isoforms identify the PCR products. (**D**) Western blot probing for MBNL1 and 2 in RBPMS knockdown and overexpression shows the MBNL1 isoform switch at the protein level. (**E**) Schematic of the different MBNL1 protein isoforms, left. Different MBNL1 and MBNL2 isoforms were overexpressed in HEK293T cells and their effect on an A5SS event in the *NCOR2* gene assessed. Schematics of *NCOR2* splicing isoforms indicates the PCR products. Western blot were probed against FLAG to verify isoform overexpression. (**F**) MBNL1 and 2 knockdown in inducible RBPMS A or in lentiviral control (P LV) proliferative PAC1 cells to assess the dependency of the *Ncor2* A5SS event to the MBNL1 isoform switch. Western blot for MBNL1 and 2 and FLAG to confirm MBNL knockdown and RBPMS A overexpression. In all the western blots TUBULIN was used as a loading control. Statistical significance was calculated using Student’s t-test (* P < 0.05, ** P < 0.01,*** P < 0.001). (**G**) Sashimi plot of the *Lsm14b* cassette exon regulated by RBPMS. Psi values represent the mean of the Psi values calculated by rMATS analysis. (**H**) RT-PCR validation of *Lsm14b* in the RBPMS knockdown and overexpression in PAC1 cells. Schematic of the *Lsm14b* event at the top and schematic of its isoforms at the left. Peptide sequence coded by the exon is shown below the schematic. (**I**) Western blot of LSM14B in RBPMS knockdown and overexpression cells. GAPDH was used for loading control.

Another post-transcriptional regulator affected by RBPMS is LSM14B, which is involved in cytoplasmic regulation of mRNA stability and translation (Brandmann et al., 2018) but also shuttles to the nucleus (Kirli et al., 2015). RBPMSA overexpression promoted skipping of *Lsm14b* exon 6, a pattern which is also seen in tissue SMCs (Fig. 6G,H), and knockdown was also seen to affect this event (Fig 6G,H), with corresponding changes in LSM14B protein (Fig. 6I). Exon 6 contains the only predicted nuclear localization signal (RPPRRR) in LSM14B (Fig. 6H) lying between the LSM and FDF domains. This suggests that RBPMS mediated AS might prevent nuclear shuttling and function of LSM14B in mRNA turnover.

### RBPMS controls the SMC transcription factor Myocardin

Myocardin (MYOCD) is a key transcription factor in SMCs and cardiac muscle (S. Li, Wang, Wang, Richardson, & Olson, 2003). Skipping of *Myocd* exon 2a in cardiac muscle and proliferative SMCs produces a canonical mRNA encoding full length MYOCD (Fig. 7A) (Creemers, Sutherland, Oh, Barbosa, & Olson, 2006; van der Veer et al., 2013). Inclusion of exon 2a in differentiated SMCs introduces an in frame stop codon, and the N-terminally truncated Myocd isoform produced using a downstream AUG codon lacks the MEF2 interacting domain and is more potent in activating SMC-specific promoters and SMC differentiation (Creemers et al., 2006; Imamura, Long, Nanda, & Miano, 2010; van der Veer et al., 2013). Significant changes in Myocd exon 2a splicing were not detected by rMATS, but manual inspection of RNA-Seq data and RT-PCR confirmed that Myocd exon 2a is more included in differentiated than proliferative PAC1 SMCs (Fig. 7B) and its inclusion decreases upon RBPMS knockdown (Fig. 7B, Fig. S2E). A conserved 200 nt region downstream of exon 2a contains two clusters of CAC motifs (Fig. 7C), suggesting that RBPMS directly activates exon 2a splicing. To better understand the regulation of the Myocd exon 2a, we created a minigene of exon 2a and its flanking intronic regions. In transfected PAC1 cells exon 2a in the minigene was included to a basal level of ∼30% and RBPMS-A or RBPMS-B significantly increased exon 2a inclusion (Fig. 7D). The Myocd minigene was also tested in HEK293 cells. As expected, exon 2a was fully skipped in the non-smooth muscle cell line, but was highly responsive to RBPMSA and B with inclusion increasing to ∼75% and 37% respectively (Fig. 7E). To test the role of the downstream CAC clusters we mutated each cluster (CAC to CCC mutations). Mutation of cluster 1 (mCAC) had a modest effect on activation by RBPMSA, although RBPMSB activity was severely impaired. Mutation of cluster 2 impaired activity of both RBPMS isoforms, while the combined mutations abolished all activation by RBPMS (Fig. 7E). Insertion of a previously defined RBPMS site (Ube2v1 from (Farazi et al., 2014)) into the double mutant minigene restored activation by RBPMS-A, although RBPMS-B had minimal activity in this context (Fig 7G). To confirm binding of RBPMS to the CAC motifs, EMSAs and UV crosslinking were carried out with *in vitro* transcribed RNAs containing the wild-type and mutant CAC clusters (Fig. 7F). RBPMS A and B were both able to bind to the wild-type RNA at similar affinities as indicated by EMSA (Fig 7F upper panels). Binding was modestly reduced by mutation of the first cluster, more severely affected by mutation of the second cluster, and eliminated by combined mutation of both clusters. UV crosslinking of RBPMS was inefficient with only a very faint signal evident with wild type, but not mutant, probes (Fig. 7F lower panels). These data therefore indicate that RBPMS directly activates Myocd exon 2a inclusion via downstream CAC clusters.

**Figure 7.**
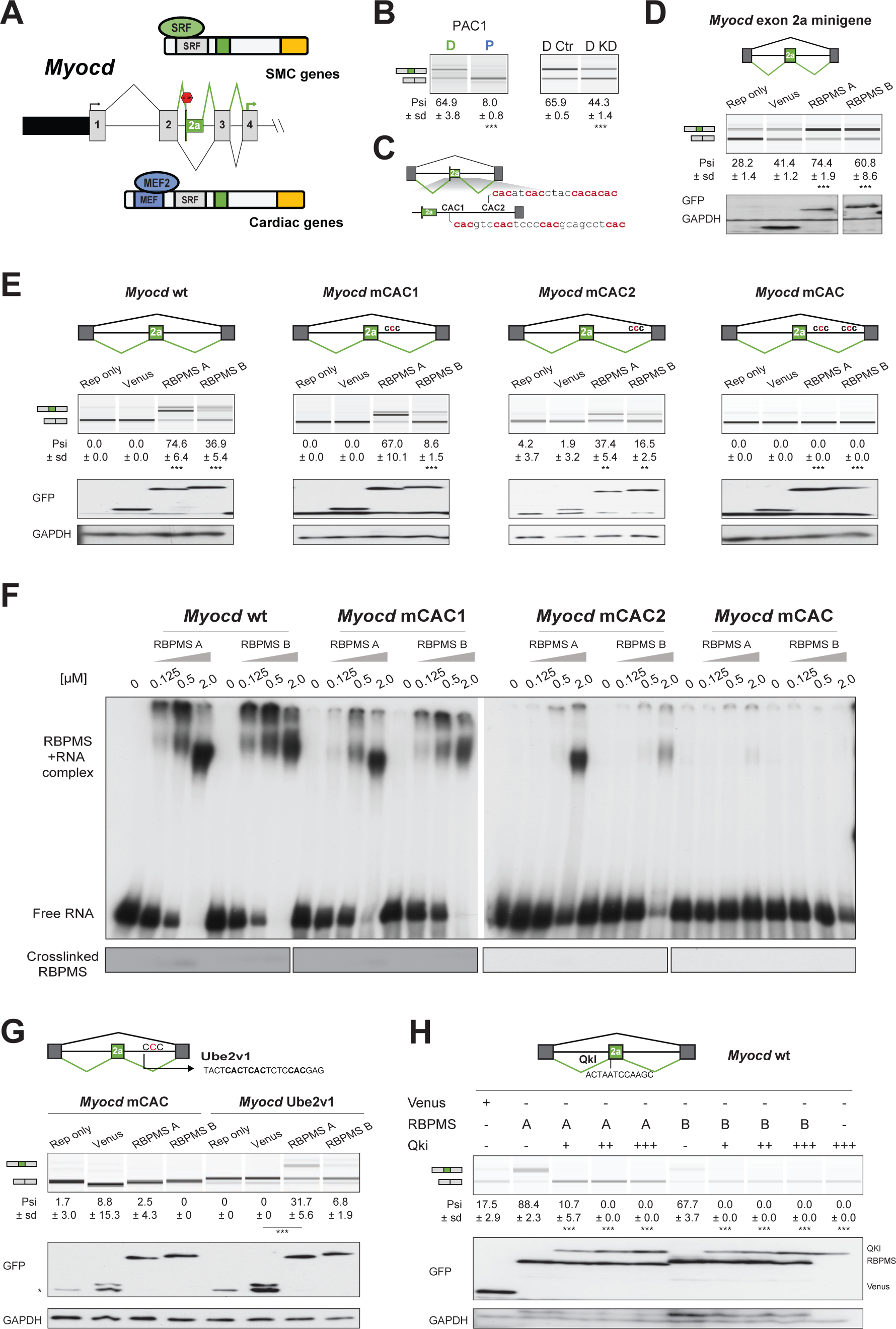
RBPMS controls splicing of the SMC transcription factor *Myocd* exon 2a. (**A**) Schematic of *Myocd* exon 2a ASE and its functional consequences upon MYOCD protein domains. (**B**) RT-PCR *Myocd* exon 2a in the PAC1 Differentiated (D) and Proliferative (P) PAC1 cells, left) and in differentiated PAC1 cells treated with control (D Ctr) or RBPMS siRNAs (D KD), right. (**C**) Schematic of the *Myocd* exon 2a splicing reporter. The two main clusters of CAC motifs, found downstream of exon 2a, are highlighted. (**D**) RT-PCRs of the effects of RBPMS A and B overexpression on *Myocd* exon 2a in PAC1 cells. Reporter only and venus controls were tested alongside. Protein expression levels are shown in the western blot probing for GFP and GAPDH as a loading control. (**E**) RT-PCR of the effects of RBPMS A and B overexpression on Myocd exon 2a splicing reporter in HEK293. RBPMS CAC sites were mutated and response to RBPMS validated by RT-PCR. Schematic of the different mutant Myocd exon 2a splicing reporters are found at the top. RBPMS overexpression was confirmed by western blot using GAPDH as a loading control. (**F**) *in vitro* binding of RBPMS A and B to Myocd exon 2a downstream intron sequence in EMSA, top, and in UV-crosslink, bottom. *in vitro* transcribed wild-type and different mCAC RNAs were incubated with RBPMS recombinant protein for binding using a serial dilution (1:4) in a range of 0.125 to 2 μM of the recombinant protein. (**G**) The UBE2V1 sequence identified to bind to RBPMS by PAR-CLIP (Farazi et al., 2014) was inserted into the mCAC Myocd reporter. Schematic of the mCAC Myocd exon 2a splicing reporter and the UBE2V1 sequence. RT-PCR of the effects of RBPMS A and B overexpression on Myocd reporter in HEK293 cells. Western blot probing for GFP and GAPDH as a loading control. (**H**) Effects of the co-expression of RBPMS and QKI, a Myocd exon 2a repressor, on Myocd exon 2a reporter in HEK293. Myocd 2a reporter schematic with QKI and potential RBPMS binding sites indicated. RBPMS and QKI overexpression was confirmed by western blot against GFP and GAPDH as a loading control. * in western blot indicate GFP product of the reporter splicing.

Myocd exon 2a is repressed by binding of the RBP QKI to the 5’ end of the exon (van der Veer et al., 2013). QKI is expressed more highly in proliferative SMCs (Llorian et al., 2016; van der Veer et al., 2013) suggesting that RBPMS and QKI could act antagonistically on AS events during phenotypic switching. To test for functional antagonism we co-transfected the Myocd minigene with the two RBPs in HEK293 cells (Fig. 7H). QKI strongly antagonized RBPMS activitation of exon 2a inclusion, even at low concentrations, showing it to be the dominant regulator (Fig 7H). Thus, splicing of Myocd exon 2a, and thereby the transcriptional activity of Myocd, is under the antagonistic control of RBPs that are preferentially expressed in differentiated (RBPMS) or proliferative (QKI) SMCs.

## Discussion

### Using super-enhancers to identify master splicing regulators

By focusing on RBPs whose expression is driven by super-enhancers (Jangi & Sharp, 2014) we identified RBPMS as a key regulator of the differentiated SMC AS program, with many of the criteria expected of a master regulator: i) it is highly up-regulated in differentiated SMCs (Fig. 1, Fig. S1). Indeed, single cell RNA-Seq identified *Rbpms* as part of a transcriptome signature of contractile mouse aorta SMCs cells (Dobnikar et al., 2018); ii) changes in RBPMS activity appear to be solely responsible for 20% of the AS changes between differentiated and proliferative PAC1 cells (Fig. 3); iii) RBPMS target splicing events are enriched in functionally coherent groups of genes affecting cell-substrate adhesion and the actin cytoskeleton, which are important for SMC cell phenotype-specific function (Fig. 5); iv) it regulates splicing and activity of other post-transcriptional regulators in SMCs (Fig. 6), and v) it regulates splicing of the key SMC transcription factor MYOCD (Fig. 7) to an isoform that promotes the contractile phenotype (van der Veer et al., 2013).

RBPMS is reported to be a transcriptional co-activator (J. Fu et al., 2015; Sun et al., 2006). However, we did not observe changes in expression levels of SMC marker genes upon RBPMS knockdown or overexpression and changes in RNA abundance were outnumbered by regulated AS events (Fig. 2). As a splicing regulator RBPMS can only affect actively transcribed genes, so it is unlikely to be sufficient to initially drive SMC differentiation. Notably, we identified RBPMS using super-enhancers mapped in adult human tissues (Hnisz et al., 2013). Combined with the ability of RBPMS to promote AS patterns characteristic of fully differentiated tissue SMCs, such as in *Cald1, Fermt2* and *Tns1* (Fig. 3F,G), this suggests that RBPMS plays a key role in maintaining a mature adult SMC phenotype. Consistent with a role in promoting a fully differentiated state, in other cell types RBPMS has anti-proliferative tumor-suppressive activity (J. Fu et al., 2015; Hou et al., 2018; Rastgoo, Pourabdollah, Abdi, Reece, & Chang, 2018).

### RBPMS is an alternative splicing regulator

RBPMS and RBPMS2 (referred to as Hermes in *Xenopus*) are present across vertebrates, while the related proteins *Drosophila* Couch Potato and *C. elegans* MEC-8 bind to similar RNA sequences (Soufari & Mackereth, 2017). RBPMS and RBPMS2 can localize to the cytoplasm and nucleus, but apart from transcriptional co-regulation (J. Fu et al., 2015; Sun et al., 2006) most attention has been paid to cytoplasmic roles in mRNA stability (Rambout et al., 2016), transport (Hornberg et al., 2013) and localization in cytoplasmic granules (Farazi et al., 2014; Furukawa, Sakamoto, & Inoue, 2015; Hornberg et al., 2013). RBPMS2 interacts with eukaryote elongation factor-2 (eEF2) in gastrointestinal SMCs, suggesting translational control (Sagnol et al., 2014). MEC-8 was reported to regulate splicing of Unc-52 in *C. elegans* (Lundquist et al., 1996), but otherwise RBPMS family members have not been reported to regulate splicing. PAR-CLIP with over-expressed RBPMS in HEK293 cells revealed its preferred binding site, but accompanying mRNA-Seq did not identify regulated AS events associated with PAR-CLIP peaks (Farazi et al., 2014). Using rMATS to re-analyse the RNA-Seq data of Farazi et al, we found a small number of regulated AS events, including the *Flnb* H1 exon (Fig. 4C-F). We readily detected specific binding of RBPMS to target RNAs by EMSA, but UV crosslinking was inefficient, even with purified protein and *in vitro* transcribed RNA (Figs. 4F,J, 7F). This suggests that CLIP might significantly under-sample authentic RBPMS binding sites. Nevertheless, the functional binding of RBPMS to *Flnb, Tpm1* and *Myocd* RNAs (Figs 4, 7), the strong enrichment of dual CAC motifs with RBPMS-regulated exons (Fig 4A), and distinct positional signatures for activation and repression of splicing (Fig. 4A,B), indicate that RBPMS acts widely to directly regulate splicing in differentiated SMCs. While earlier reports have shown that RBPMS can act in other steps in gene expression, our data provide the strongest evidence to date for a widespread molecular function of RBPMS as a splicing regulator.

Cell-specific splicing programs are usually driven by more than one regulatory RBPs. A number of splicing regulatory RBPs are known to promote the proliferative SMC phenotype, including QKI, PTBP1, and SRSF1 (Llorian et al., 2016; van der Veer et al., 2013; Xie et al., 2017). The extent to which these proteins coordinately regulate the same target ASEs remains to be established. RBPMS and QKI antagonistically regulate at least two targets in addition to *Myocd* (Fig. 7). The *Flnb* H1 exon is activated by RBPMS (Fig. S2, Fig. 4) and repressed by QKI (J. Li et al., 2018), while the penultimate exon of *Smtn* is repressed by RBPMS (Fig. S2E) but activated by QKI (Llorian et al., 2016). Moreover, QKI binding motifs are significantly associated with exons regulated during SMC dedifferentiation, and with exons directly regulated by RBPMS overexpression or knockdown (data not shown). It is therefore an interesting possibility that RBPMS and QKI might target a common set of ASEs perhaps acting as antagonistic master regulators of differentiated and proliferative SMC phenotypes. The two RBPs show reciprocal regulation of expression levels during SMC dedifferentiation (Fig. S1D), which combined with antagonistic activities could lead to switch like changes in many ASEs. For Myocd splicing, QKI appears to have dominant activity, driving skipping of exon 2a even when RBPMS is present at higher levels (Fig. 7G), so full inclusion of exon 2a requires the presence of RBPMS and absence of QKI. The logic of this regulatory input can explain inclusion of Myocd exon 2a in SMC and skipping in cardiac muscle and is consistent with the observation that RBPMS is super-enhancer associated both in vascular SMCs and heart left ventricle, while QKI is super-enhancer associated in left ventricle but not differentiated SMCs. Notably, QKI also controls a significant fraction of ASEs regulated during myogenic differentiation of skeletal muscle cells, but it promotes differentiated myotube AS patterns (Hall et al., 2013), in contrast to its promotion of dedifferentiated splicing patterns in SMCs.

### Activity of RBPMS isoforms

*RBPMS* was originally characterized as a human gene that encoded multiple isoforms of an RBP (Shimamoto et al., 1996), but differential activity of common RBPMS isoforms has not previously been reported. Human RBPMSA and B have similar activity for co-regulation of AP1 transcriptional activity (J. Fu et al., 2015) although a third human-specific isoform had lower activity. We found differential activity of RBPMSA and B upon some ASEs (Figs. 4,7). In general, RBPMSA has higher activity particularly for repressed targets (Fig. 4F, Fig. S4A, B). The differential activity was seen with similar levels of overall expression, although we could not rule out the possibility of variation in nuclear levels as GFP-tagged RBPMS was predominantly cytoplasmic. Nevertheless, some ASEs were differentially responsive to RBPMSA or B while others responded equally (e.g. compare *MPRIP* with *ACTN1*, Fig. S4A). The differential activity upon *Tpm1* splicing could not be accounted for by differences in RNA binding by RBPMSA or B (Fig. 4F,H). Therefore, it is possible that RBPMS repressive function depends on other interactions mediated by the 20 amino acid RBPMSA C-terminal (Fig 1C). While the RRM domain is sufficient for RNA binding and dimerization *in vitro* (Sagnol et al., 2014; Teplova et al., 2016), previous studies have shown the extended C terminal region downstream of the RRM domain to be involved in several aspects of RBPMS/RBPMS2 function including granular localization in retinal ganglion cells (Hornberg et al., 2013) and interaction with cFos in HEK293 cells (J. Fu et al., 2015). *In vitro* assays suggested that the C-terminal region increases RNA binding affinity and possibly the oligomeric state of RBPMSA (Farazi et al., 2014) and the C-terminal 34 amino acids of *Xenopus* RBPMS2 is required for binding to *Nanos1* RNA *in vivo* (Aguero et al., 2016). In preliminary studies we have also found the C-terminal region to be essential for splicing regulation (data not shown). Future work will aim to address the mechanisms of RBPMS splicing activation and repression as well as isoform-specific differential activity.

### RBPMS targets numerous mRNAs important for SMC function

SMC phenotypic switching involves interconversion between a contractile phenotype and a motile, secretory, proliferative state (Owens et al., 2004). The actin cytoskeleton and its connections to the extracellular matrix (ECM) via focal adhesions are central to the function of both cell states, but with contrasting outcomes: tissue-wide contraction or independent movement of individual cells. RBPMS-mediated AS plays a major role in remodelling the actin cytoskeleton, the integrin adhesome (Horton et al., 2015) and ECM components in the two cell states. This is reflected in the similar GO terms shared by RBPMS regulated AS events and the entire PAC1 cell AS program (Fig. 5, Fig. S5). The importance of many of the RBPMS target genes to SMC function is further indicated by their proximity to super-enhancers in SMC tissues (Fig. 5B,C). Indeed, three targets - ACTN1, FLNB and TNS1 - are all super-enhancer associated, interact directly with both actin and integrins and are components of distinct axes of the consensus integin adhesome (Horton et al., 2015). ACTN1 is a major hub in the network of proteins affected by RBPMS (Fig. 5C). The AS event in ACTN1 produces a functional Ca^2+^-binding domain in the motile isoform, but lack of Ca^2+^ binding in the differentiated isoform stabilizes ACTN1 containing structures in contractile cells (Waites et al., 1992). Many other RBPMS regulated AS events have not previously been characterized. Focal adhesion complexes and the integrin adhesome are mechanosensitive complexes that connect the cytoskeleton and ECM, and are hubs of regulatory tyrosine phosphorylation signalling. We found a small network of RBPMS regulated AS events in the receptor tyrosine phosphatase PTPRF (Fig 2E) and two interacting proteins PPFIA1 and PPFIBP1 (Fig. S2E). Another focal adhesion associated target is PIEZO1 (Fig. 2E), a mechanosensitive ion channel that is important in SMCs during arterial remodelling in hypertension (Retailleau et al., 2015). An RBPMS repressed exon lies within the conserved Piezo domain immediately adjacent to the mechanosensing “beam” (Liang & Howard, 2018). ECM components affected by RBPMS include fibronectin (FN1), which interacts directly with integrins, and HSPG2 (Fig. S2E). Notably, HSPG2 (also known as Perlecan) is the basement membrane heparan sulfate proteoglycan that is the identified splicing target of MEC-8 in *C. elegans* (Lundquist et al., 1996).

In addition to direct regulation of numerous functionally related targets, by directly targeting transcriptional and post-transcriptional regulators RBPMS has the potential for more widespread action (Figs. 6,7). RBPMS regulated events in MBNL1 and 2 modulate their splicing regulatory activity (Sznajder et al., 2016; Tabaglio et al., 2018; Tran et al., 2011). Changes in secondary AS targets are challenging to observe in a short duration overexpression experiment. Nevertheless, the modest change in *NCOR2* splicing appears to be attributable to an RBPMS-induced switch to less active MBNL isoforms (Fig 6F). The regulated event in LSM14B (Fig 6G-I) also has the potential to regulate mRNA stability, or an as yet uncharacterized nuclear role of LSM14B (Kirli et al., 2015).

We also identified the SMC transcription factor as a direct target of RBPMS (Fig. 7). A similar role has been shown for the proposed myogenic AS master regulator RBM24 (Jangi & Sharp, 2014), which stabilizes mRNA of the transcription factor Myogenin by binding to its 3’UTR (Jin, Hidaka, Shirai, & Morisaki, 2010). Similarly, by activating inclusion of *Myocd* exon 2a (Fig. 7), RBPMS directs production of a Myocardin isoform that more potently promotes the differentiated SMC phenotype (van der Veer et al., 2013). Additional effects upon transcription could also be conferred by RBPMS activation of the FLNB H1 exon (Fig. S2E, Fig. 4). FLNB is primarily an actin binding and adhesion protein, but inclusion of the H1 hinge domain allows nuclear localization and antagonism of the transcription factor FOXC1 in epithelial cells (J. Li et al., 2018). FOXC1 and FOXC2 are expressed at higher levels in adult arteries than any other human tissue (Consortium, 2013). The modulation by RBPMS of FLNB isoforms therefore provides another route for indirect transcriptome regulation. The importance of Filamin RNA processing in SMCs by adenosine to inosine editing of FLNA was also recently highlighted by the cardiovascular phenotypes arising from disruption of this editing (Jain et al., 2018). A number of splicing regulators also influence miRNA processing (Michlewski & Caceres, 2019), so it is an interesting possibility that RBPMS might affect processing of SMC miRNAs such as miR143-145 (Boettger et al., 2009; Cordes et al., 2009).

RBPMS2 plays an important role in SMCs of the digestive tract. The *RBPMS2* gene is associated with super-enhancers in stomach smooth muscle (Supplementary File 1) and is expressed early in visceral SMC development and at lower levels in mature cells (Notarnicola et al., 2012). Ectopic RBPMS2 overexpression led to loss of differentiated contractile function *via* translational upregulation of *Noggin* (Notarnicola et al., 2012; Sagnol et al., 2014). This contrasts with our observations that RBPMS exclusively promotes differentiated SMC AS patterns. RBPMS2 is expressed at low levels in PAC1 and primary aorta SMCs (Fig 1) and its knockdown was without effect. Nevertheless, ectopic expression of RBPMS2 in PAC1 or HEK293 cells promoted differentiated SMC AS patterns in a similar manner to RBPMSA (Fig. 4, Fig. S4), suggesting that RBPMSA and RBPMS2 have intrinsically similar molecular activities. The reasons for the apparent discrepancy between the promotion by RBPMS2 of differentiated SMC splicing patterns, but de-differentiated visceral SMC phenotypes, remain to be resolved. Possible explanations include variations in cell-specific signalling pathways, pre-mRNA and mRNA targets, interacting protein partners, post-translational modifications and subcellular localization, all of which could differentially modulate RBPMS and RBPMS2 activity in different SMC types.

In conclusion, our data vindicate the proposal that tissue-specific AS master regulators might be identified by the association of their genes with superenhancers (Jangi & Sharp, 2014), paving the way for the identification of further such regulators in other tissues. While our data suggest that RBPMS has a critical role in SMCs, it is likely to play important roles in other cell types where its expression is also super-enhancer driven, including cardiac muscle and embryonic stem cells. Our approach aimed to identify AS master regulators common to diverse smooth muscle types (vascular, bladder, stomach). However, SMCs show a great deal of diversity (Fisher, 2010), even within single blood vessels (Cheung, Bernardo, Trotter, Pedersen, & Sinha, 2012). The splicing regulator Tra2 β is responsible for some splicing differences between tonic and phasic SMCs (Shukla & Fisher, 2008), and it is possible that other RBPs might act as master regulators of some of these specialized SMC types. Our future studies aim to understand the mechanisms of splicing regulation by RBPMS, the role of the RBPMS regulated splicing program in controlling different aspects of SMC phenotype and the potential role of subversion of this program in cardiovascular diseases.

## Materials and Methods

### Identification of potential master AS regulators

Locations of human super-enhancers (genome build Hg19) were taken from the data sets UCSD_Aorta, UCSD_Bladder, BI_Stomach_Smooth_Muscle and BI_Skeletal_Muscle in (Hnisz et al., 2013). Associated genes were obtained using the UCSD Table Browser (GREAT version 3.0.0) with the Association rule: “Basal+extension: 5000 bp upstream, 1000 bp downstream, 1000000 bp max extension, curated regulatory domains included”. We used the 1542 human RBPs from (Gerstberger et al., 2014) to identify potential master AS regulators within each set of super-enhancer proximal genes (Supplementary File 1).

### DNA constructs

Coding sequences of rat *Rbpms* isoforms were PCR amplified from differentiated PAC1 cell cDNA and cloned into XhoI/EcoRI sites of the pEGFP-C1 vector (Clontech) and into EcoRI/XhoI sites of the pCI-neo-3x-FLAG vector (Rideau et al., 2006) to generate N-terminal Venus and 3xFLAG tagged *in vivo* overexpression constructs. The two major *Rbpms* isoforms identified were RBPMS A (XM_006253240.2/ XP_006253302.1) and RBPMS B (NM_001271244.1/ NP_001258173.1). QKI construct has been described in a previous study (Llorian et al., 2016).

Splicing reporters of *Tpm1* exon 3 and *Actn1* exon NM and SM were described in (Gooding et al., 2013; Gromak et al., 2003). *Myocd* exon 2a and *Flnb* exon H1 splicing reporters were obtained by PCR amplification of the target exons and respective flanking intron regions from genomic PAC1 DNA, approximately 250 bp upstream and downstream for *Myocd* and 500 bp for *Flnb*. PCR products were subsequently cloned into XhoI/EcoRV and NotI/SphI sites of pCAGGs-EGFP vector, which contains a GFP expression cassette with an intron inserted into its second codon (Wollerton, Gooding, Wagner, Garcia-Blanco, & Smith, 2004). Point mutations of the CAC motifs in the splicing reporters were generated by PCR using oligonucleotides that contained A to C mutations. Intronic regions containing CACs from *Tpm1, Flnb* and *Myocd* were PCR amplified and cloned into HindIII/EcoRI sites of pGEM4Z (Promega) for *in vitro* transcription. DNA constructs were confirmed by sequencing. All the oligonucleotides used for cloning and mutagenesis are found in Supplementary File 6.

### Cell culture, transfection and inducible lentiviral cell line

Rat PAC1 pulmonary artery SMCs (Rothman et al., 1992) were grown to a more differentiated or proliferative state as described in (Llorian et al., 2016). HEK293T cells were cultured following standard procedures. *Rbpms* siRNA mediated knockdown in PAC1 cells was performed as in (Llorian et al., 2016). Briefly, 10^5^ differentiated PAC1 cells were seeded in a 6 well plate. After 24 hours, cells were transfected using oligofectamine reagent (Invitrogen) and 90 pmols of Stealth siRNAs from Thermo Fisher Scientific (siRNA1: RSS363828, GGCGGCAAAGCCGAGAAGGAGAACA). A second treatment was performed after 24 hours using lipofectamine2000 (Thermo Fisher) and siRNA at the same concentration of the first treatment. Total RNA and protein were harvested 48 hours after the second knockdown. C2 scrambled siRNA was used as a control in the knockdown experiments (Dharmacon, C2 custom siRNA, AAGGUCCGGCUCCCCCAAAUG). For *Mbnl1* and *Mbnl2* siRNA knockdown, the *Mbnl1* THH2 siRNA (CACGGAAUGUAAAUUUGCAUU) and *Mbnl2* specific siRNA (Dharmacon, GAAGAGUAAUUGCCUGCUUUU) were used (Gooding et al., 2013). 3xFLAG N-terminally tagged rat RBPMSA cDNA was cloned into pInducer22 (Meerbrey et al., 2011) using the Gateway system. Generation of stable PAC1 cell lines with pInducer22 vector only (LV) or pInducer22-RBPMSA was done as follows. Lentiviral particles were produced in HEK293T cells by transient transfection using 30 µl of Mirus TransIT-lenti (MIR6604), 7 µg 3xFLAG tagged RBPMSA cDNA and 0.75 µg of the packaging plasmids gag, pol, tat & VSV-G transfecting 2 × 10^6^ cells per 10 cm dish. After 24 hours the medium was transferred to 4°C and replaced with fresh medium. After a further 24 hours the medium was removed, pooled with the first batch, spun at 1000 g for 5 minutes and filtered through 0.45 micron PVDF filter. Lentiviral particles were diluted 1:2 with fresh DMEM medium containing Glutamax, 10% FBS and 16 µg/ml polybrene and used to replace the medium on PAC1 cells plated 24 hours earlier at 10^4^/35mm well, setting up 2 wells for pInducer22 vector only and 6 wells for RBPMSA. Fresh medium was added 24 hours later and the populations amplified as necessary. To induce expression of 3xFLAG RBPMSA the cells were plated at 4 × 10^5^ cells per 35 mm well ± 1µg/ml doxycycline harvesting RNA and protein 24 hours later.

For transient transfections of HEK293 cells with splicing reporters and effectors, lipofectamine 2000 reagent was used and cells harvested 48 hours after transfection. To verify knockdown and transfection efficiency, total cell lysates were obtained by directly adding protein loading buffer to the cells. Lysates were run on a SDS-PAGE, followed by western blot against RBPMS and loading controls. See Supplementary File 6 for information on the antibodies used in this study. To monitor changes in splicing and mRNA abundance, RNA was extracted using TRI reagent (Sigma) according to manufacturer’s instructions, DNase treated with Turbo DNA-free kit (Thermo Fisher) and cDNA synthesized, as described below, followed by PCR and QIAxcel or qRT-PCR analysis.

### qRT-PCR and RT-PCR

cDNA was prepared using 1 μg total RNA, oligo(dT) or gene-specific oligonucleotides and SuperScript II (Life technologies) or AMV RT (Promega) as described in manufacturer’s protocol. qRT-PCR reactions were prepared with 50 ng of cDNA, oligonucleotides for detection of mRNA abundance and SYBER Green JumpStart Taq Ready Mix (Sigma). Three-step protocol runs were carried out in a Rotor-Gene Q instrument (QIAGEN). Analysis was performed in the Rotor-Gene Q Series Software 1.7 using the Comparative Quantitative analysis. To normalize the relative expression values, two housekeeper genes were included in each experiment (*Gapdh* and *Rpl32*) and their geometric mean used for normalization. Expression values were acquired from biological triplicates.

PCRs with 50 ng of the prepared cDNAs were carried out to detect the different mRNA splicing isoforms of the reporters using the oligonucleotides detailed in Supplementary File 6. For visualization and quantification of the PSI values, PCR products were resolved in a Qiaxcel Advanced System (QIAGEN) and PSI calculated within the QIAxcel ScreenGel software. A minus RT cDNA, representative of each triplicate, and a no template PCR reactions were also included in all the experiments (data not shown). Statistical significance was tested as previously described for gene expression (paired Student t-test for lentiviral experiments). PSI values are shown as mean (%) ± standard deviation (sd). For lentiviral RBPMSA overexpression, ΔPSI values determined by RT-PCR were derived from at least three independent transductions, and no events responded to doxycycline treatment in cells transduced with the empty pINDUCER22 vector (Figure S2D). Statistical significance was tested by a two-tailed Student’s t test, paired for lentiviral experiments and unpaired for all the others (* P< 0.05, ** P<0.01, *** P<0.001).

### Immunostaining

For immunodetection of RBPMS in PAC1 cells, differentiated and proliferative cells were grown on coverslips, fixed with 4% paraformaldehyde (PFA) for 5 minutes, rinsed with phosphate buffer saline (PBS) and permeabilised with 0.5% NP-40 for 2 minutes followed by PBS washes. Coverslips were incubated with blocking buffer (1% BSA in PBS) for 1 hour and incubated with RBPMS primary antibody diluted in blocking buffer for another hour. Coverslips were rinsed and secondary antibody in blocking buffer applied to the coverslips which were incubated for 1 hour. Coverslips were washed and mounted on ProLong Diamond Antifade with DAPI (Thermo Fisher Scientific). All the steps were carried out at room temperature. Images were acquired from a fluorescence microscope (Zeiss Ax10, 40X) attached to CCD AxioCam and analysed on AxioVision (v4.8.2).

### RNAseq analyses

Total RNA from three biological replicates of RBPMS knockdown in differentiated PAC1 and three populations of RBPMS inducible overexpression in proliferative PAC1 cells were extracted for RNAseq. Cells were lysed with TRI-reagent and total RNA purified by Direct-zol purification column (Zymo Research) followed by DNase treatment. RNAseq libraries of polyA selected RNAs were prepared with NEBNext ultra II RNA library prep kit for Illumina. Both RNA and RNAseq libraries were checked for their qualities. Barcoded RNAseq libraries were then multiplexed across two lanes of an Illumina HiSeq4000 platform for sequencing on a 150-bp paired-end mode, providing around 60 million reads per sample.

To investigate AS changes in vascular smooth muscle dedifferentiation, rat aortas were isolated from 8-12 weeks old Wistar rats. Aortas were briefly treated for 30 minutes at 37°C with 3 mg/ml collagenase (Sigma C-0130) to help in cleaning away the adventitia. The tissue was finely chopped and either used directly to make tissue RNA or enzymatically dispersed to single cells. This was achieved by treating the tissue pieces with 5 ml 1 mg/ml elastase (Worthington Biochemical Corporation LS002292) for 30 minutes at 37°C and then 5 ml collagenase added for a further 1-2 hours. Cells were washed & counted and plated at 4 × 10^5^ cells/ml in M199 media containing 10%FBS, 2 mM Glutamine and 100 U/ml Penicillin-Streptomycin in a suitable dish according to the cell number. To promote the proliferative state, SMCs were 1:2 passaged switching to DMEM media containing Glutamax and 10% FBS and harvested at passage 9. For RNAseq, total RNA was harvested from three replicas, each a pool of 5 rats, from rat aorta tissue (T), enzymatically dispersed single cultured SMCs (SC), passage 0 (P0) and passage 9 (P9). Total RNA extraction was carried out with Tri-reagent (Sigma). Libraries for mRNAseq were prepared using Ribozero and TrueSeq kits and sequencing performed on a HiSeq2000 platform in a paired-end mode.

### mRNA abundance analysis

Read trimming and adapter removal were performed using Trimmomatic version 0.36 (Bolger, Lohse, & Usadel, 2014). Reads were aligned using STAR version 2.5.2a (Dobin et al., 2013) to the Rat genome Rnor_6.0 release-89 obtained from Ensembl and RSEM package version 1.2.31 (B. Li & Dewey, 2011) was used to obtain gene level counts. mRNA abundance analysis, was carried out with DESeq2 package (version 1.18.1) (Love et al., 2014) within R version 3.4.1 (https://www.r-project.org/). Genes were considered to be differential expressed with p-adj less than 0.05 in the paired analysis (Supplementary File 2).

### AS changes analysis

rMATS v3.2.5 (Shen et al., 2014) was used for detection of differential alternative splicing. rMATS analysis was carried out allowing for new splicing event discovery using the flag novelSS 1. rMATS calculates the inclusion levels of the alternative spliced exons and classifies them into five categories of AS events (Fig. S2B): skipped exons (SE), mutually exclusive exons (MXE), alternative 5′ and 3′ splice sites (A5SS and A3SS) and retained intron (RI). Before further analysis, the results from reads on target and junction counts were filtered to include only events with a total of read counts above 50 across the triplicates in at least one of the conditions compared. Removal of events with low counts discarded false positive events. Only ASE with an FDR less than 0.05 were considered significant and a minimal inclusion level difference of 10% imposed to significant AS events. Finally, to identify specific AS events, unique IDs were created using the AS type, gene name and the chromosomal coordinates of the regulated and flanking exons (Supplementary File 3).

For visualization of differentially spliced exons, sashimi plots were generated using rmats2sashimiplot (Gohr & Irimia, 2018). The sashimi plots show the RNAseq coverage reads mapping to the exon-exon junctions and Psi values from rMATS. Twenty eight ASE identified by rMATS in the RBPMS knockdown or overexpression were also validated by RT-PCR in the same manner as described in the qRT-PCR and RT-PCR section. ΔPsi predicted from the RNAseq analysis and the ΔPSI observed in the RT-PCR were then tested for a Pearson correlation in RStudio (http://www.rstudio.com/).

For comparison and visualization of the overlap between the genes with different mRNA abundance and the genes differentially spliced in the RBPMS knockdown, RBPMS overexpression and the PAC1 dedifferentiation, proportional Venn diagrams were made using BioVenn (Hulsen, de Vlieg, & Alkema, 2008). Venn diagrams were also generated for the visualization of the common AS events across RBPMS knockdown, overexpression and the PAC1 or aorta tissue dedifferentiation datasets.

### RBPMS motif enrichment analysis

RBPMS motif enrichment analyses in the regulated cassette exons (SE) of RBPMS knockdown and overexpression, PAC1 and Aorta tissue dedifferentiation were performed using the toolkit MATT (Gohr and Irimia, 2018). Cassette exons in transcripts identified by rMATS with significant changes (FDR < 0.05 and |ΔPSI| > 10%) were used to test enrichment or depletion of CACN_1-12_CAC RBPMS recognition element (Farazi et al., 2014), against a background set of unregulated exons defined as events with FDR > 0.1 and |ΔPSI| < 5%. 250 bp of the flanking intronic regions were examined for RBPMS motif signals. The motif enrichment scores were first obtained using the test_regexp_enrich command of the Matt suite in the quant mode with statistical significance determined using a permutation test with 50,000 iterations. This module inherently divides the examined regions (exons or 250bp of the introns) into thirds and provides positional information for the occurrence of the RBPMS motif.

RNA maps for distribution of the RBPMS motif were generated using the rna_maps command in the Matt suite. Here the unregulated set of exons was randomly downsampled to include a final background of 2000 events only. Additionally, the program was instructed to scan only 135 bp of the cassette exon and 250 bp at either end of the flanking introns. A sliding window of 31 was used for scanning the motif and the statistically significant regions (p < 0.05) for enrichment or depletion were identified by the permutation tests specifying 1000 iterations.

### Gene ontology and PPI analysis

Enrichment for gene ontology terms in the differentially abundant and spliced genes were obtained from Gorilla (Eden, Navon, Steinfeld, Lipson, & Yakhini, 2009). Two unranked lists of genes, target and background lists, were used for the GO analysis. The target list contained either the significant differential abundant genes (p-adj < 0.05 and log2 Fold Change greater than 1 or less than −1) or the significant differentially spliced genes (FDR < 0.05 and ΔPsi threshold of 10%). Background gene lists were created for the PAC1 experiments and aorta tissue by selecting the genes whose expression were higher than 1 TPM in either of the conditions analyzed. For visualization, only the top 5 enriched GO terms of each category (biological process, cell component and molecular function) were shown in Figure 5A and Figure S5. The complete list of enriched terms in differentially spliced and abundant genes can be found in Tables S4 and S5.

A Protein-protein interaction network for the genes differentially spliced by RBPMS was constructed using the STRING v10.5 database (Szklarczyk et al., 2017). RBPMS regulated genes were obtained by merging two lists: i) overlap of genes concordantly differentially spliced in the RBPMS knockdown and PAC1 experiments and, ii) overlap of genes concordantly differentially spliced in the RBPMSA overexpression and the aorta tissue datasets that were shown to be regulated in the same range in both conditions. A cut off of ΔPsi greater than 10% was also applied to the RBPMS regulated gene list, similar to the GO analysis. The human database was chosen for the analysis and the following parameters applied to the PPI network: confidence as the meaning of the network edges, experiments and database as the interaction sources and high confidence (0.700) as the minimum required interaction score. STRING functional enrichments, using the whole genome as statistical background, were also included for visualization. Human super-enhancer associated genes from Supplementary File 1 were highlighted.

### Recombinant protein

Rat RBPMS A and B with a 3xFLAG N terminal tag were cloned into the BamHI/XhoI sites of the expression vector pET21d, for expression of recombinant RBPMS containing a T7 N-terminal tag and a His_6_ C-terminal tag in *E. coli*. Recombinant RBPMS A protein was purified using Blue Sepharose 6 and HisTrap HP columns whereas RBPMS B was purified only through the latter, since low binding was observed to Blue Sepharose 6. The identity of purified recombinant proteins was confirmed by western blot (Fig. S4G) and mass mapping by mass spectrometry.

### *In vitro* transcription and binding

α^32^ P-UTP labelled RNA probes were *in vitro* transcribed using SP6 RNA polymerase. For EMSAs, a titration of the recombinant RBPMS A and B (0, 0.125, 0.5 and 2 μM) was incubated with 10 fmol of *in vitro* transcribed RNA in binding buffer (10 mM Hepes pH 7.2, 3 mM MgCl_2_, 5% glycerol, 1 mM DTT, 40 mM KCl) for 25 minutes at 30°C. After incubation, samples were run on a 4% polyacrylamide gel. For UV-crosslinking experiments, the same binding incubation was performed followed by UV-crosslink on ice in a Stratalinker with 1920 mJ. Binding reactions were then incubated with RNase A1 and T1 at 0.28 mg/ml and 0.8 U/ml respectively, for 10 minutes at 37°C. Prior to loading the samples into a 20% denaturing polyacrylamide gel, SDS buffer was added to the samples which were then heated for 5 minutes at 90°C.

### Statistical analysis

Analysis and quantification of RNAseq and RT-PCR experiments were described in their respective sections with further information of the tests used in the different experiments present in the figure legends. Graphics were generated in RStudio (http://www.rstudio.com/).

### Data availability

mRNAseq of RBPMS (knockdown and overexpression) and Aorta tissue dedifferentiation data from this study have been deposited in xxxxx, accession number xxxxxx.

## Acknowledgements

We thank Elisa Monzon-Casanova for helpful comments on the manuscript, and Juan Mata, Sushma Grellscheid, Vasudev Kumanduri and Elisa Monzon-Casanova for help and advice on RNA-Seq data analysis.

## Author contributions

Conceptualization, C.W.J.S.; Investigation, E.E.N.S., C.G., M.L., A.G.J., F.R., A.B.; Writing – Original Draft, E.E.N.S. and C.W.J.S.; Writing – Review & Editing, C.G. A.G.J., M.L., S.S.; Funding Acquisition, C.W.J.S., S.S.

## Supplementary Figure Legends

**Figure S1.**
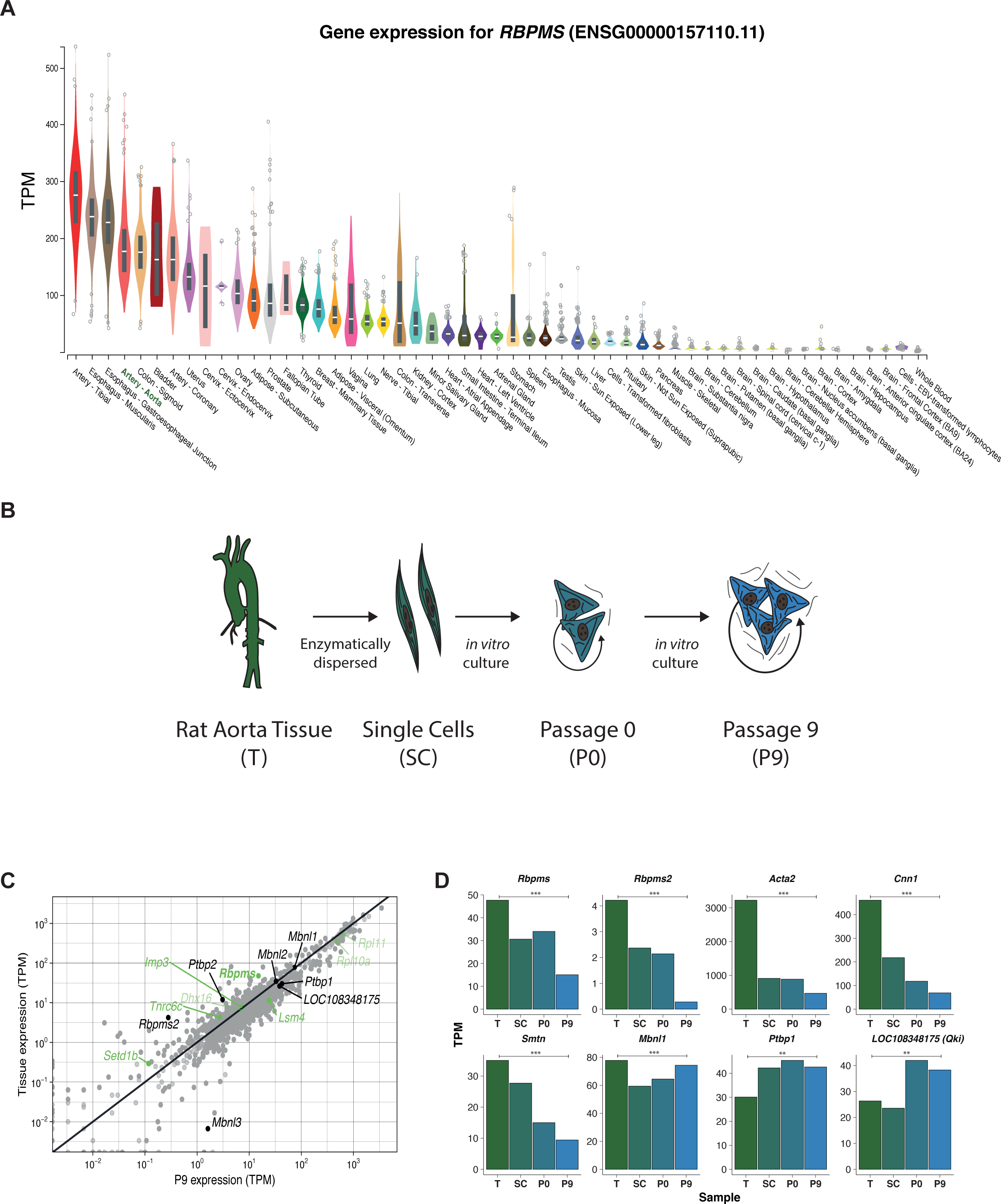
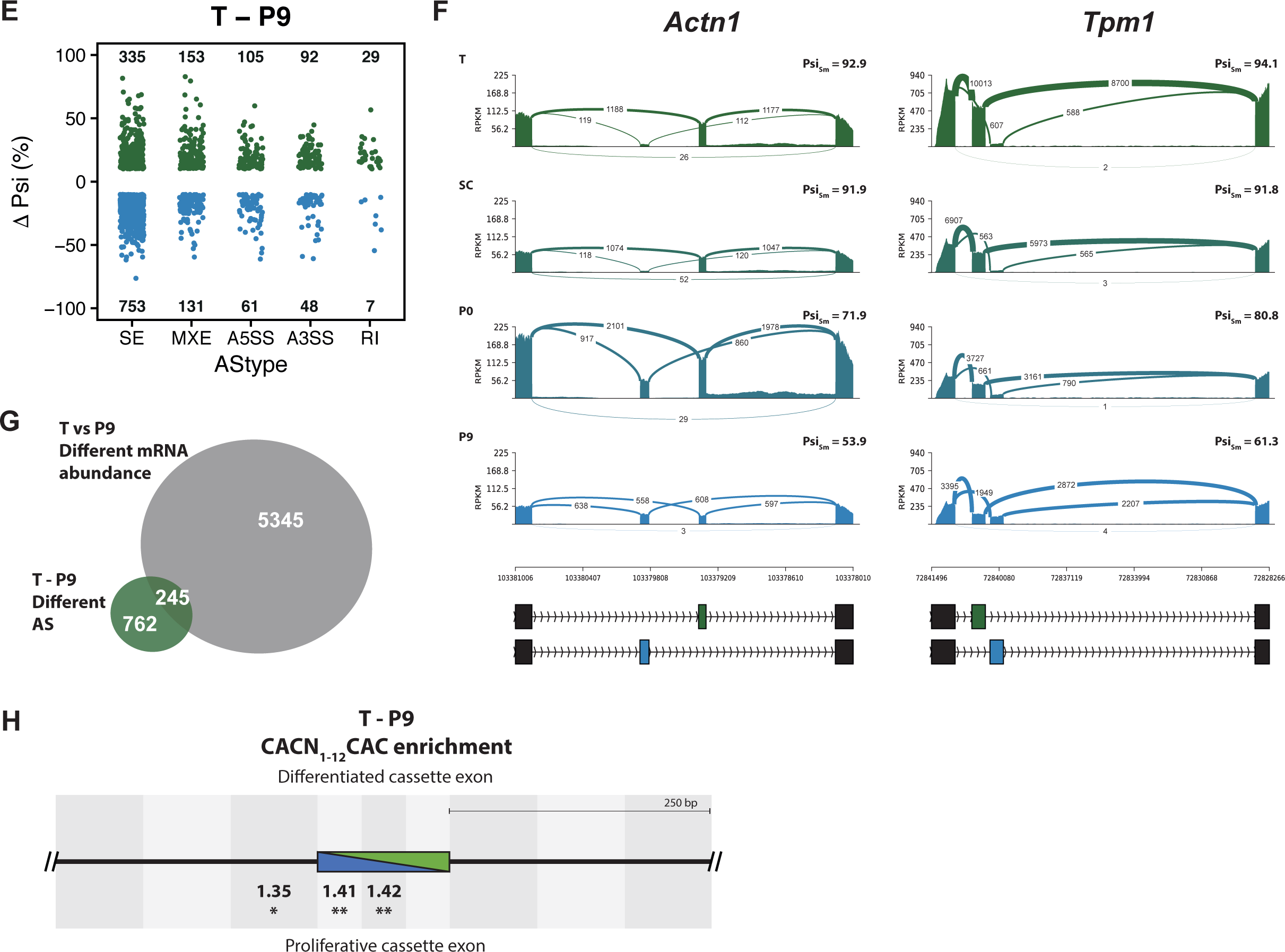
(**A**) *RBPMS* mRNA expression across different human tissues in the GTEX database (Consortium, 2013). (**B**) Schematic of the experimental design of the rat aorta tissue dedifferentiation. (**C**) mRNA abundance of RBP genes in differentiated aorta tissue (T) and proliferative passage 9 (P9). RBPs whose genes were found associated with smooth muscle tissue super-enhancers (Fig. 1B) are highlighted in green. Additionally, other well characterized RBPs and Rbpms2 are labelled in black. LOC108348175 is the Ensembl annotated rat Qki like gene. Statistical significance in mRNA abundance changes between T and P9 was calculated by DESeq2 and is indicated in dark and light gray color (padj < 0.05 and non-significant changes respectively). mRNA levels are expressed in transcripts per million (TPM). Cpsf4l gene was not found in this dataset. (**D**) mRNA abundance changes measured in transcripts per million (TPM) during rat aorta dedifferentiation (T, SC, P0 and P9). Genes shown are *Rbpms* and its paralog *Rbpms2*, SMC differentiation markers, *Acta2, Cnn1* and *Smtn*, and other RBPs, *Mbnl1, Ptbp1* and *Qk1*. (**E**) AS changes when comparing T and P9 conditions. Only significant ASE with FDR < 0.05 and Δpsi greater than 10% are shown. ASE were classified as skipped exons (SE), mutually exclusive exons (MXE), 5’and 3’ alternative splice sites (A5SS and A3SS) and retained intron (RI). The number of events differentially alternatively spliced are shown in the graph. (**F**) Sashimi plot of the SMC mutually exclusive splicing markers, *Actn1* and *Tpm1*, in rat aorta tissue dedifferentiation. Psi values represent the mean of the Psi values calculated by rMATS. (**G**) Overlap of the number of genes affected at the mRNA abundance and AS in the T and P9 comparison. (**H**) RBPMS motif enrichment in differentially alternatively spliced SE in the T and P9 comparison. A pair of CAC separated by 1 to 12 nt was used as the RBPMS motif. Values indicate the motif enrichment. Statistical significance was calculated by the Matt tool (* P < 0.05, ** P < 0.01,*** P < 0.001).

**Figure S2.**
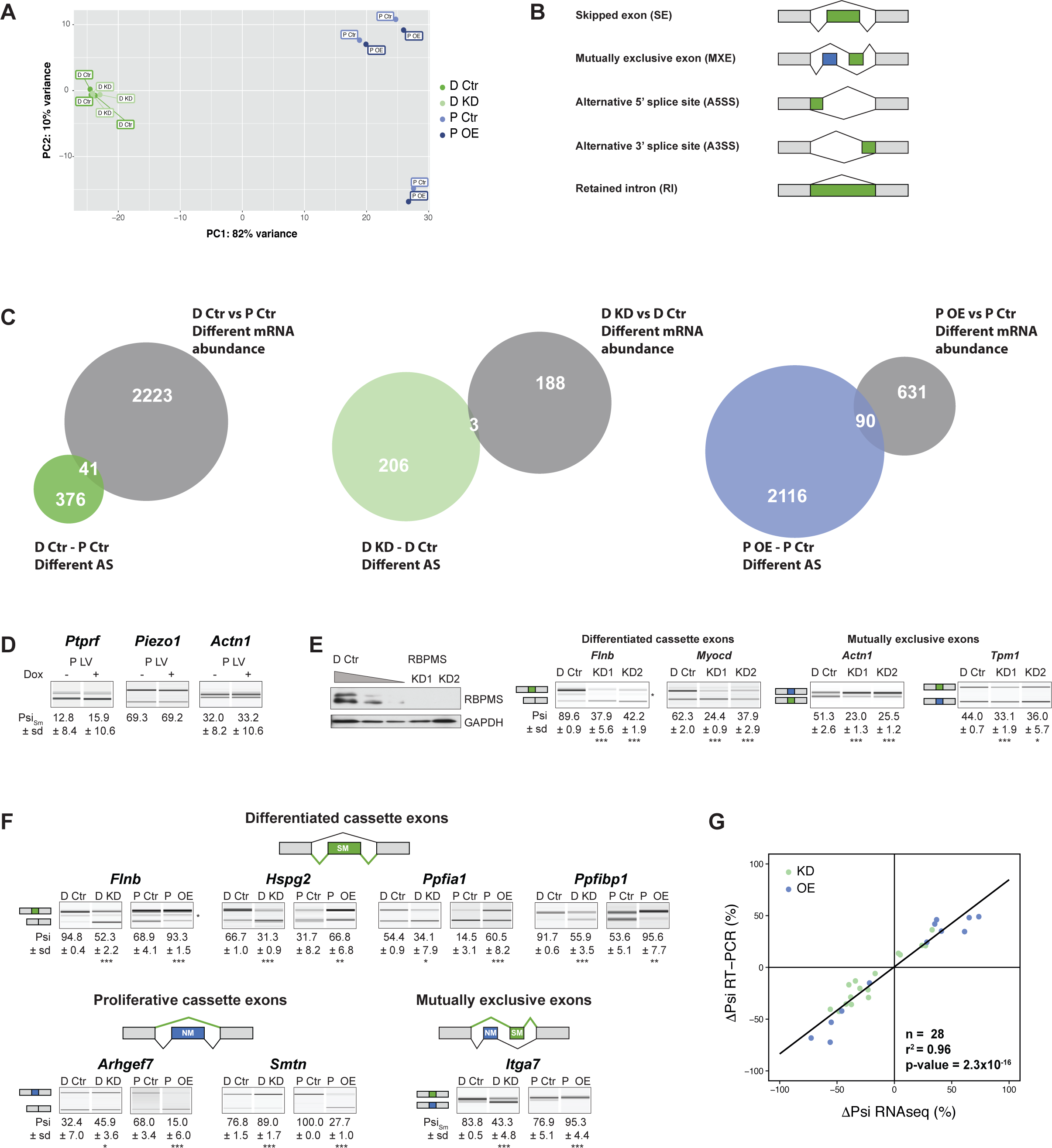
(**A**) Principal Component Analysis (PCA) based upon mRNA abundance variance of RBPMS knockdown (D Ctr and D KD) and overexpression (P Ctr and P OE) replicates. (**B**) Schematics of the different AS types detected by rMATS. (**C**) Overlap of the number of genes affected at the mRNA abundance and AS in the PAC1 dedifferentiation and RBPMS knockdown and overexpression. (**D**) Validation of RBPMS regulated ASE in lentiviral control populations (P LV) upon doxycycline treatment. (**E**) Validation of a second RBPMS siRNA (KD2, Stealth siRNAs, Thermo Fisher Scientific (RSS363829): CGCUUCGAUCCUGAAAUCCCGCAAA). Western blot probing for RBPMS in differentiated PAC1 cells treated with one of the siRNA. GAPDH was used as a loading control in the western blot. RNAs were extracted and RT-PCR carried out to validate the ASE characterized before for KD1. (**F**) Validation of differentiated and proliferative cassette and mutually exclusive splicing events by RT-PCR for both RBPMS knockdown and overexpression. Schematics indicates the splicing isoform corresponding to the PCR product. In (**D**), (**E**) and (**F**), values shown are the mean ± sd of Psi (n = 3). Statistical significance was calculated using Student’s t-test (* P < 0.05, ** P < 0.01,*** P < 0.001). (**G**) Psi correlation between the estimated ΔPsi from the rMATS analysis of the RBPMS knockdown and overexpression RNAseq experiments and the observed ΔPsi from the RT-PCR validations of RBPMS experiments (n=28). Black line indicates the linear regression model. For statistical significance, a Pearson correlation test was carried out in RStudio and the results are shown in the right corner of the plot. r_2_ is the correlation coefficient and the p-value indicates the statistical significance of the correlation.

**Figure S3.**
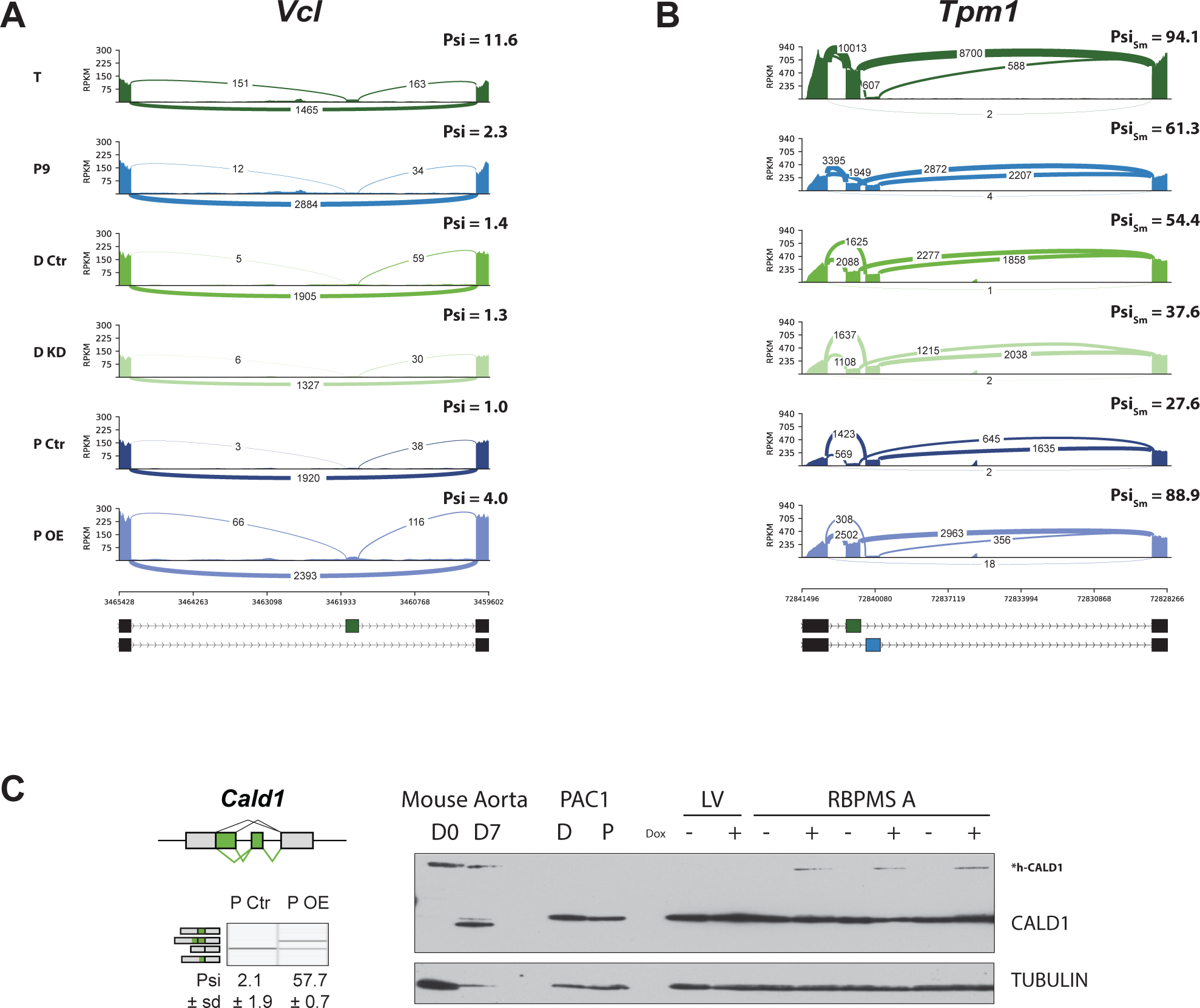
(**A**) Sashimi plot of the *Vcl* cassette exon regulated during aorta tissue dedifferentiation (FDR < 4×10_-14_) and by RBPMS overexpression (FDR < 1.2×10_-12_). Psi values represent the mean of the Psi values calculated by rMATS analysis. (**B**) Sashimi plot of the *Tpm1* mutually exclusive exon regulated during aorta tissue dedifferentiation and by RBPMS overexpression. Psi values represent the mean of the Psi values calculated by rMATS analysis for the SM exon 2. (**C**) RT-PCR validation of *Cald1* ASE (A5SS and SE) in the RBPMS overexpression in proliferative PAC1 cells. Schematic of the *Cald1* ASE and *Cald1* isoforms amplified in the PCR are shown on the top and left, respectively. The switch in *Cald1* isoform was shown at the protein level by western blot probing for CALD1. The specificity of the isoform switch was also confirmed in *Mus musculus* differentiated (D0) and proliferative (D7) SMC samples from aorta tissue. The larger smooth muscle specific isoform of CALD1 is indicated by h-CALD1. Values shown are mean ± sd (n = 3) of percentage of exon inclusion. Statistical significance was calculated using Student’s t-test (* P < 0.05, ** P < 0.01,*** P < 0.001).

**Figure S4.**
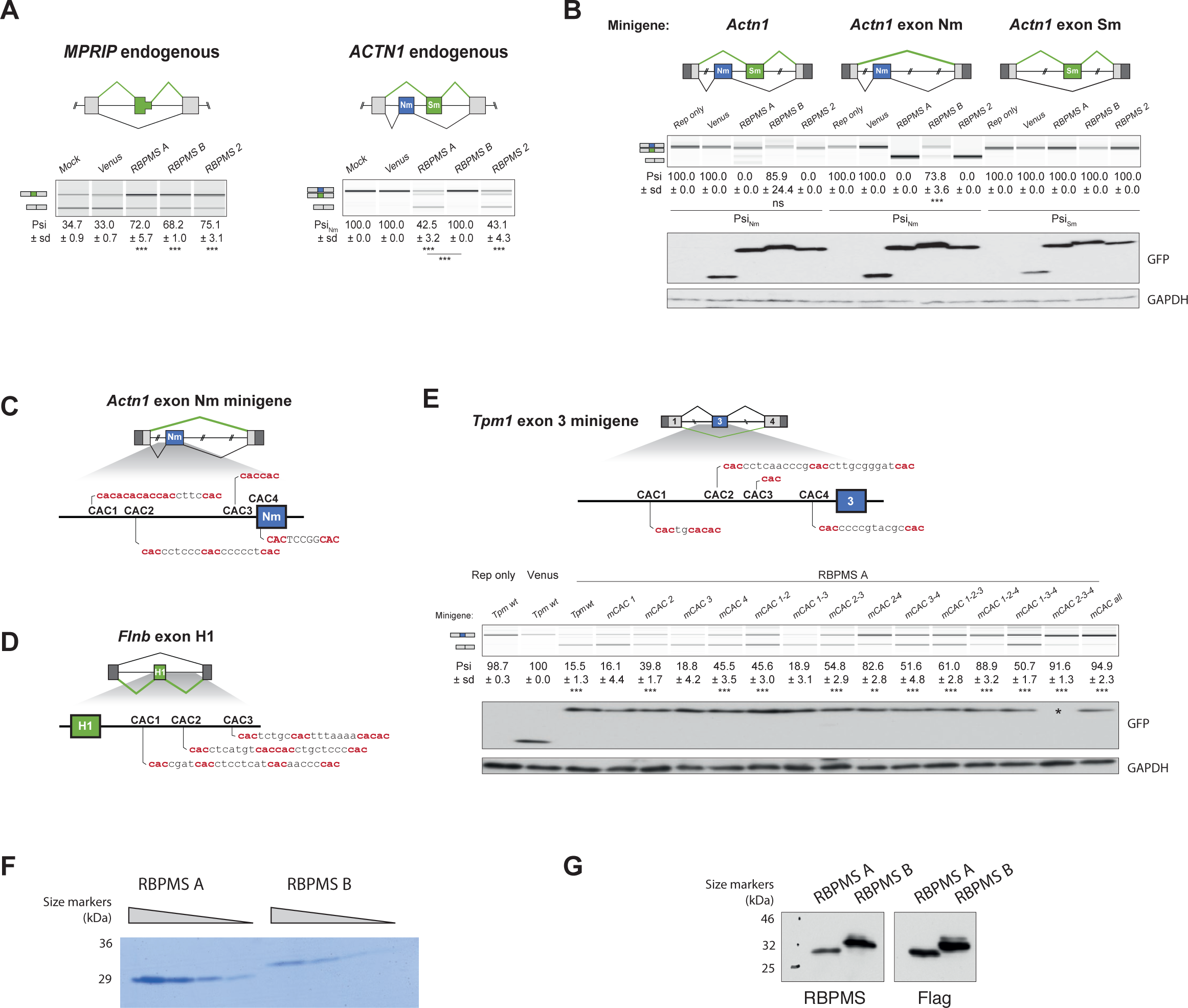
(**A**) RT-PCR validation of endogenous *MPRIP* and *ACTN1* ASE in the RBPMS isoforms and RBPMS2 overexpression in HEK293 cells. Schematics of *MPRIP* SE and *ACTN1* MXE are found at the top. (**B**) *Actn1* splicing upon RBPMS overexpression in HEK293 cells. *Actn1* reporters containing both the Sm and Nm exons or only the Sm or Nm exon were tested upon RBPMS overexpression. Schematics of the different *Actn1* splicing reporters tested are shown at the top. Western blot to verify RBPMS overexpression were probed for GFP and GAPDH as a loading control. (**C**) Schematic of potential RBPMS CAC motifs upstream of *Actn1* exon Nm and (**D**) downstream of *Flnb* exon H1. (**E**) Mapping RBPMS cis elements in the *Tpm1* exon 3 splicing reporter. Potential RBPMS CAC motifs upstream of exon 3 are highlighted in the schematic at the top. Combinations of different CAC mutations were tested for RBPMS A overexpression in HEK293 cells. overexpression was assessed by western blot probing for GFP and GAPDH as a loading control. * No overexpression was detected in this lane although the Psi differs from the reporter only Psi. In (**A**), (**B**) and (**E**), schematics of the splicing isoforms identify the PCR products and values shown are mean ± sd (n = 3) of Psi. In case of MXE the Sm exon inclusion is shown. Statistical significance was calculated using Student’s t-test (* P < 0.05, ** P < 0.01,*** P < 0.001). (**F**) RBPMS A and B recombinant proteins. Purified recombinant RBPMS A and B were analyzed in a 20% denaturing polyacrylamide gel. A BSA titration was run in parallel for quantification of the recombinant protein. (**G**) Recombinant protein sequences were confirmed by western blot probing for RBPMS and FLAG. Full western blot images are shown.

**Figure S5.**
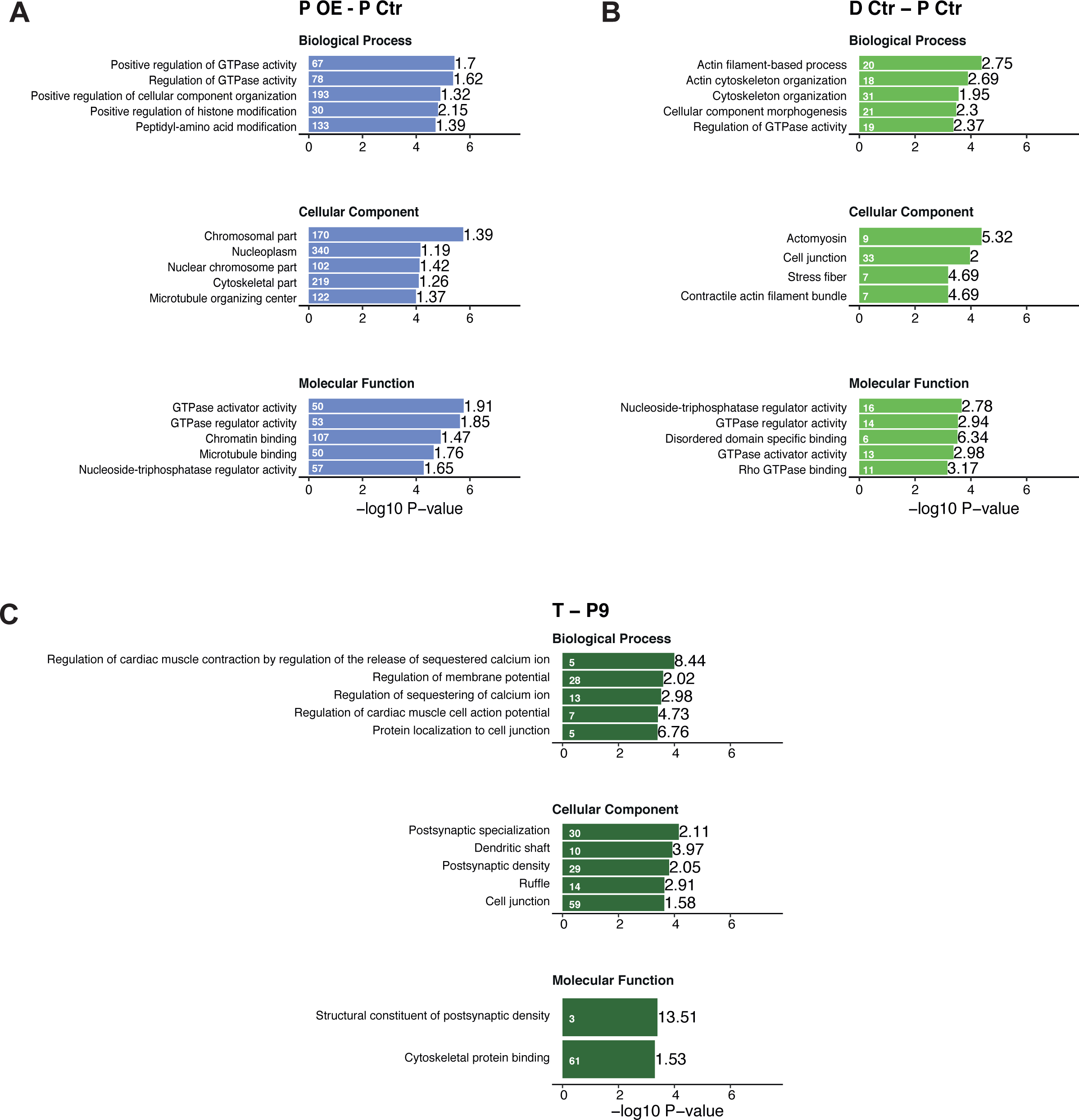
Top five enriched GO terms from Gorilla GO analysis of differentially spliced genes. (**A**) RBPMS overexpression in proliferative PAC1 cells. (**B**) PAC1 dedifferentiation. (**C**) Rat aorta tissue dedifferentiation. Values within and in front of the bars indicate the number of genes in the enriched term and the enrichment relative to the background list.

## Supplementary Data Files

**Supplementary File 1.** RBP genes associated with super-enhancers in human tissues.

**Supplementary File 2.** Genes with significant changes in mRNA abundance in aorta dedifferentiation (T vs P9), PAC1 dedifferentiation (D Ctr vs P Ctr), RBPMS knockdown (D KD vs D Ctr) and RBPMS overexpression (P OE vs P Ctr).

**Supplementary File 3.** Genes with significant changes in mRNA splicing in aorta dedifferentiation (T - P9), PAC1 dedifferentiation (D Ctr - P Ctr), RBPMS knockdown (D KD - D Ctr) and RBPMS overexpression (P OE - P Ctr).

**Supplementary File 4.** GO terms significantly enriched in the genes differentially spliced in aorta dedifferentiation (T - P9), PAC1 dedifferentiation (D Ctr - P Ctr), RBPMS knockdown (D KD - D Ctr) and RBPMS overexpression (P OE - P Ctr).

**Supplementary File 5.** GO terms significantly enriched in the genes with differential mRNA abundance.

**Supplementary File 6.** Oligonucleotides and antibodies used in this study.

